# Variation and process of life history evolution in insular dwarfism as revealed by a natural experiment

**DOI:** 10.1101/2020.12.23.424186

**Authors:** Shoji Hayashi, Mugino O. Kubo, Marcelo R. Sánchez-Villagra, Hiroyuki Taruno, Masako Izawa, Tsunehiro Shiroma, Takayoshi Nakano, Masaki Fujita

## Abstract

Islands are a classic focus for evolutionary studies. One subject of much interest has been the evolution of “dwarfs”, significantly smaller island mammals relative to their continental counterparts. Although a consensus has been achieved that multivariate ecological causes are behind body size changes, the processes involved remain largely unexplored. Life history variables, including the age of first reproduction, growth rate, and longevity, are probably key to understanding the process of insular dwarfism. The Japanese Archipelago with numerous islands offers a unique natural experiment of evolution into different sizes within the same group of organisms, deer. Thus, we investigated eight deer populations with a total number of 52 individuals exhibiting body size variation, both extant and fossil, to clarify the effect of insularity on life history traits. We applied several methods to both extant and extinct populations to resolve life history changes among deer populations. Skeletochronology, using lines of arrested growth formed in long bones (femur and tibia), successfully reconstructed body growth curves and revealed a gradual change in growth trajectories reflecting the degree of insularity. Slower growth rates with prolonged growth periods in more isolated deer populations were revealed. An extensive examination of bone microstructure further corroborated this fact, with much slower growth and later somatic maturity evident in fossil insular deer isolated for more than 1.5 Myr. Finally, mortality patterns assessed by demographic analysis showed variation among deer populations, with a life history of insular populations shifting toward the “slow life”.

## Introduction

Among the most iconic examples of visible evolutionary changes are island mammals that dramatically changed in body size [1-3]. While elephants, hippos, and deer on islands across the world became small, rodents and other small mammals became larger [2]. These changes are which led to the suggestion of the “island rule” [1]. Classic examples are found in fossil species: the pygmy elephant (*Elephas falconeri*) of Sicily, which has a body mass only 1% of its mainland ancestor [4], and a gigantic gymnure (*Deinogalerix koenigswaldi*) of Miocene Gargano having a body mass about 200 times heavier than its ancestor [5]. This leads to a hypothesis of the optimal body size of mammals on isolated islands, although the particular optimum of an insular population varies with the characteristics of the islands and the species, and with interactions among species [5, 6].

Moreover, since body size has a close relationship with various aspects of organisms’ life, recent studies have focused on the relationship between body size and life history, including growth rate, age of reproduction, and longevity on islands [7-9]. A reduction in the body size of large mammals has also been associated with a modification of the growth trajectory [10, 11], which in turn affects the lifespan, e.g., the timing of sexual maturity or longevity. Life history theory, which extended from the *r*/*K* selection theory in ecology [12], proposed the idea that mammalian populations can be placed along a fast-slow continuum [13]. Species that adapted to unstable environments, under which high intrinsic rates of natural increase (*r*) were adaptive, evolved a set of life history traits, such as rapid body growth, early sexual maturity, many offspring for a clutch, and short life span, occupying the “fast” end of the continuum (formerly referred as “*r*-selected”). On the other hand, in stable environments, the populations are maintained near their carrying capacity (*K*), where resultant intraspecific competitions favor those which have slow body growth, late sexual maturity, a small number of offspring for a clutch, and a long life span, occupying the “slow” end of the continuum (“*K*-selected”). Based on this theoretical background, life history traits were estimated for fossil insular species. Such studies have been conducted by applying histological analysis of bones developed to reconstruct the paleoecology of dinosaurs and other Mesozoic vertebrates [7, 14-16], the method being further recognized by mammalian paleontologists as a useful tool to infer life history traits in mammals [8, 17-22]. Köhler and Moyà-Solà [8] showed that in a fossil dwarfed bovine *Myotragus balearicus* from the Balearic Islands of the Mediterranean, its long bone histology showed lamellar-zonal tissue throughout the cortex, which is characteristic of ectothermic reptiles. The bone microstructures indicate that *M. balearicus* is estimated to have had a low metabolism and a very slow growth rate. This life history shift toward a slow life in *M. balearicus* was further corroborated by analyses of tooth enamel microstructure, tooth eruption and mortality pattern, which implied delayed somatic maturity, lower mortality and extended longevity of this species [9, 23, 24]. Followed these paleohistological analyses were fossil mammals from other Mediterranean islands: e.g. a fossil cervid (*Candiacervus*) from Crete showing a slight shift toward a slower life history [18, 25], and a fossil dwarf hippo (*Hippopotamus minor*) from Cyprus with no evident modification in bone histology [19]. The bone histological observations and life history implications of fossil dwarfs have been accumulated in the last decade, nonetheless, these studies were based on different species from single islands, therefore it was hard to generalize the evolution of body size and life history on islands. Multiple comparisons of closely related species or populations from both mainlands and islands are crucial to drawing general conclusions.

Further important issues awaiting to be explored are the process of body size and life history evolution [26], including how ecological factors affect processes, such as the lack of predators and competitors, resource limitation on islands, founder effects of island immigrants, and the remoteness and duration of isolation [2, 5, 6, 27, 28]. Drastic body size changes have been reported for fossil species or populations [2].

Nevertheless, as far as we focused only on fossil evidence, ecological processes undergone during these changes remain unclear because of the lack of detailed ecological information. To achieve this change, again, studies on the mainland and insular populations of a single extant species are particularly useful, as they provide ecological contrasts between focal populations and allow confounding factors other than living on islands to be controlled. Variation in isolation durations among the populations can therefore mimic the process of insular settlement.

In this study, we focused on deer in the Japanese Archipelago, both extant and extinct populations from both islands and the mainland (Fig. 1). Our targets cover an extant sika deer (*Cervus nippon*) population that was introduced recently (c.a. 400 years ago) on a small island, four extant insular and mainland deer populations [sika deer, Reeves’s muntjac (*Muntiacus reevesi*)] with variable body sizes, and extinct insular deer populations (*Cervus astylodon* and Muntiacini gen. et. sp. indet., referred to as “the Ryukyu muntjac”) that were isolated for more than 1.5 Myr on the Okinawa Island [29], representing different degrees of insular effect. We also included an extinct gigantic mainland deer (*Sinomegaceros yabei*) from the Late Pleistocene deposits on the Honshu mainland, having a comparable body size to the largest extant deer; the moose (*Alces alces*) [30]. The immense variation of the island’s size and duration of isolation observed in the eight understudied deer populations enabled us to infer the process of body size and life history changes on islands.

**FIGURE 1.**
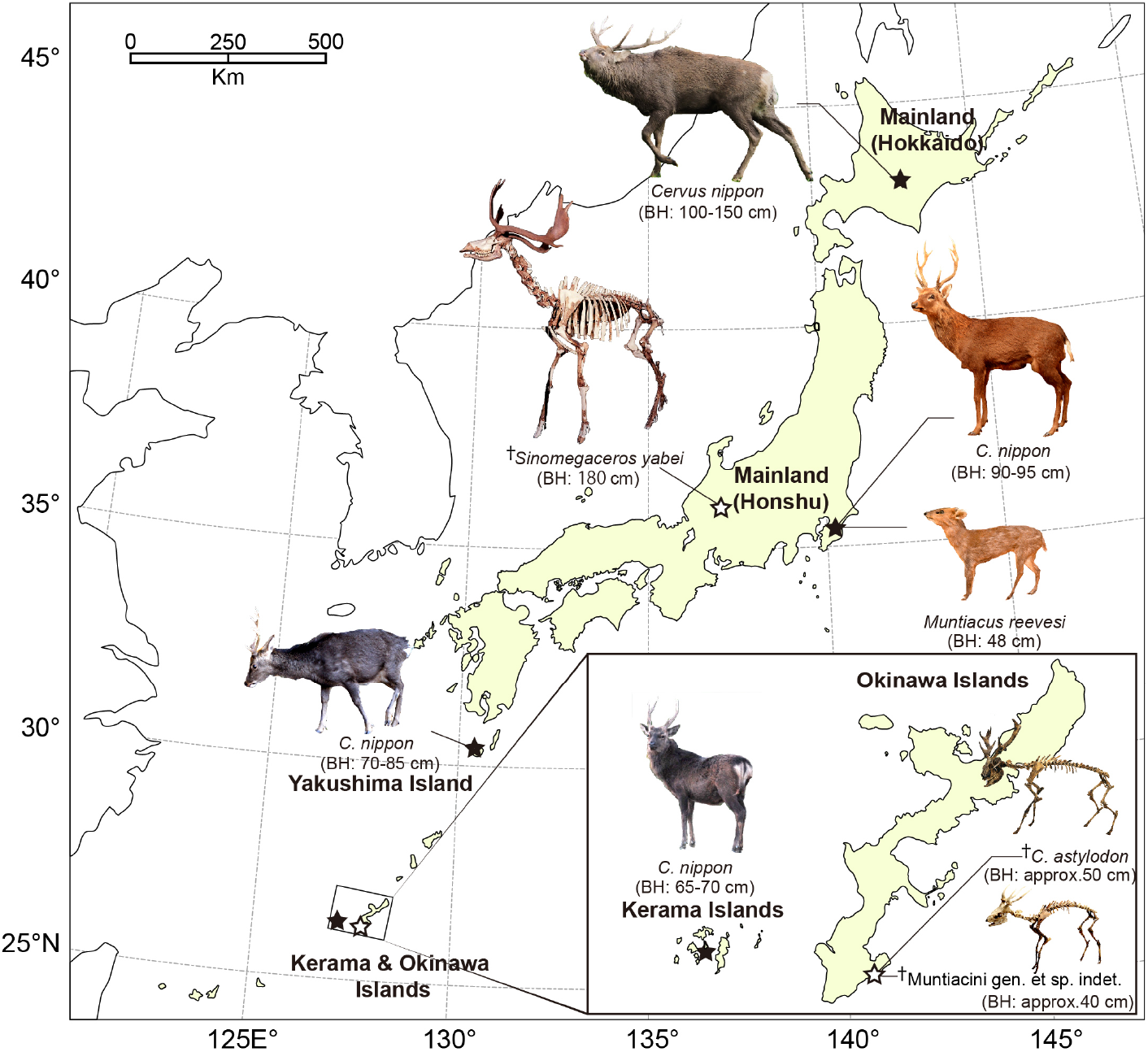
Locations of the sampled deer populations in Japan. Dagger (^†^) indicates fossils excavated from Late Pleistocene deposits. Localities of extant taxa are shown in closed stars, whereas those of extinct taxa are in open stars. The approximate body height (BH) was also indicated for each species/population. Photography of fossil cervids was courtesy of the Gunma Museum of Natural History (*Sinomegaceros yabei*) and Okinawa Prefectural Museum & Art Museum (*Cervus astylodon* and Muntiacini gen. et sp. indet.).

This study aims to test whether the different degree of insular effect results in the change of life history traits among Japanese deer populations. Specifically, we hypothesize that a gradual change in life history traits of deer populations is expected according to the intensity of insular effects, with longer isolation on smaller islands, resulting in a shift toward the more pronounced slow life. For this objective, we first applied skeletochronology to long bones, using lines of arrested growth (LAGs) within the primary periosteal bone tissue, to depict body growth rates. Obtained growth curve parameters were statistically compared among deer populations to test the hypothesis. Subsequently, detailed qualitative descriptions of bone histology were conducted aiming to corroborate the skeletochronological findings on growth patterns and somatic maturity. We focused on periosteal bone tissue and vascularization pattern as an indicator of the relative bone growth rate [8, 31]. Lastly, we conducted demographic analyses of age data, which were collected either through ecological surveys or analysis of tooth wear obtained from fossil deer assemblages, to quantitatively compare mortality patterns among deer populations.

## Materials and Methods

### 1. Locality and specimen information for the sampled deer used in the histological analyses

For histological investigation and estimation of growth curves from LAGs, we used femora and tibiae of extant and fossil deer. All extant deer were wild individuals and their skeletal specimens were stored in museums. Both tibia and femur were sampled from the same individual for the extant deer. On the other hand, all fossil specimens were excavated in disassociated conditions. Therefore, we considered that each of the femora or tibiae originated from a different individual. A summary of the sampled populations together with the sample size is provided in Table 1.

**TABLE 1.**
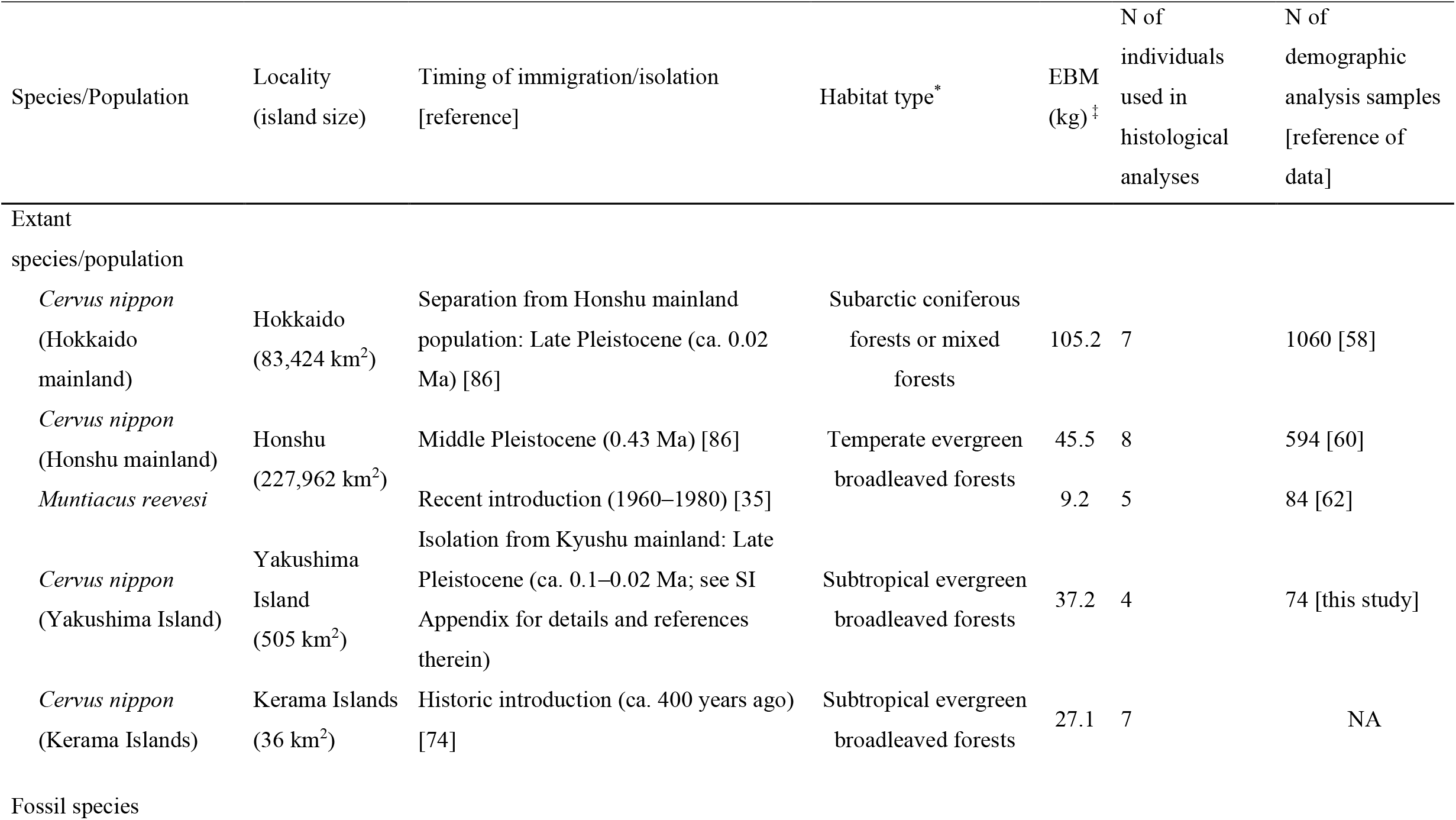
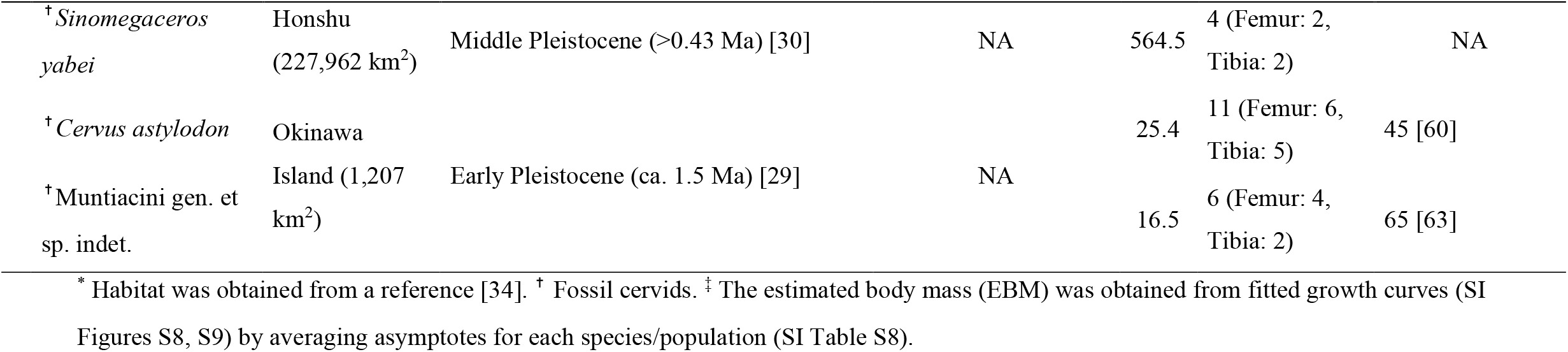
Summary of the general information of sampled Japanese deer populations.

#### 1-1. Extant deer populations

Sika deer from Hokkaido mainland: all individuals (N = 7) were hunted in Hokkaido Prefecture (43°N, 143°E), northern Japan (Fig. 1). Hokkaido is one of the four main islands constituting the Japanese archipelago and was considered “mainland” for this study. Hokkaido sika deer inhabit subarctic coniferous forests mixed with cool temperate deciduous trees and are known to be the largest subspecies (*C. nippon yesoensis* [32-34]). The complete skeletons are housed in the Hokkaido University Museum (HOUMVC), Sapporo, Japan. The age at death was recorded in the museum inventory list, which was assessed based on tooth eruption for fawns and yearlings and the tooth cementum annuli formed at the roots of the incisors for adults.

Sika deer (N = 8) and Reeves’s muntjac (N = 5) from Honshu mainland: all individuals were hunted or collected in Chiba Prefecture (35°N, 140°E, Fig. 1), which is in the central region of Honshu. Honshu is one of the four main islands constituting the Japanese archipelago and was considered “mainland” for this study. Both of these species inhabit evergreen broadleaved forests in the temperate zone. Reeves’s muntjac was introduced to Chiba Prefecture ca. 1960–1980 following an escape from a private park [35]. The source population is unknown, but the body size of these deer is smaller than the continental population but closer to that of the Taiwanese population [35]. It is not clear whether Reeves’s muntjac studied here or those in Taiwan are dwarfed populations compared with their counterpart in the continent. The complete skeletons of these individuals are stored at the Natural History Museum and Institute, Chiba (CBM), Japan. The age at death was recorded in the museum inventory list, which was assessed based on tooth eruption or the cementum annuli for the sika deer and from tooth eruption and the molar wear condition for Reeves’s muntjac.

Sika deer from Yakushima Island: all individuals (N = 4) were hunted or collected on Yakushima Island (30°18′N, 130°30′E, Fig. 1) in Kagoshima Prefecture, Japan, where they inhabit subtropical evergreen broadleaved forests. Yakushima Island deer are considered to be the subspecies *C. nippon yakushimae* [33] and exhibit considerable phenotypic differences from the mainland populations, such as small body size, simplified antlers, and shortened limbs [33, 36]. Yakushima deer diverged from the population on Kyushu Island (one of the four main islands constituting the Japanese archipelago) ca. 0.1 Mya when the Ohsumi Strait was formed between the two islands [37]. Yakushima Island was possibly connected to Kyushu Island by a land bridge during the Last Glacial Maximum (ca. 25,000 ya), but the genetic differentiation between the two populations implies a much longer period of isolation for Yakushima deer [38, 39]. The complete skeletons are stored at Tochigi Prefectural Museum (TPM), Utsunomiya, Japan. The age at death was recorded in the museum inventory list, which was assessed based on tooth eruption for fawns and yearlings and the tooth cementum annuli of the incisors and molars for adults.

Sika deer from Kerama Islands: all individuals (N = 7) were collected as carcasses resulting from natural deaths on Kerama Islands (26°12′N, 127°24′E, Fig. 1) in Okinawa Prefecture, Japan, by two of the authors (MI and TS) during their fieldwork. Sika deer were introduced to Kerama Islands from Kyushu ca. 400 years ago possibly from Kagoshima Prefecture, Kyushu mainland, Japan [40], and inhabit subtropical evergreen broadleaved forests. These deer exhibit the phenotypic characteristics of insular dwarf deer (similar to Yakushima Island deer) having smaller body size than the possible source population of Kagoshima [40] and so are also regarded as a subspecies of sika deer (*C. nippon keramae*) [33]. Most individuals were complete skeletons, but some bone elements were lost at the time of collection in the field for some individuals. All of the specimens are stored at the Faculty of Science, University of the Ryukyus (URB), Nishihara, Japan. The age at death was recorded in the inventory list, which was assessed based on tooth eruption for fawns and yearlings and the tooth cementum annuli for adults. However, no reliable age at death was obtained for an individual URB-MAM-34.

#### 1-2. Fossil deer populations

*Cervus astylodon* (femur N = 6, tibia N = 5) and Muntiacini gen. et sp. indet. (the Ryukyu muntjac, femur N = 4, tibia N = 2): both of these fossil deer species were excavated from the Hananda-Gama Cave site (26°8′7″N, 127°45′32″E, Fig. 1) in the southern part of Okinawa Island [41]. These animal remains are mainly isolated materials consisting of limb bones and teeth and are considered to have fallen into the cave through fissures or sinkholes following the decomposition of the carcasses [41]. The detailed geological age of these fossils is not clear due to a lack of suitable specimens for radiocarbon dating. However, based on radiocarbon dating at other sites that have yielded fossils of these species, the geological age of both species has been estimated as the Latest Pleistocene (>20 ka, [41]). *C. astylodon* has been recognized as a typical insular dwarf due to its very small body size and shortened limbs [2]. The Ryukyu muntjac used to be referred to as *Dicrocerus* sp. [2], which is a genus that is known mostly from Miocene Europe. However, the finding that the antlers have a clear burr and grow from the posterior parts of the frontals oriented backward clearly indicates that it does not belong to this genus (Gertrud Rößner, personal communication to MOK). Rather, it appears to be closely related to *Muntiacus* based on the presence of tusk-like canines and a lack of antlers in the females, neither of which is found in *Dicrocerus* [42]. A full revision of this species is required, including a detailed description and comparisons with other fossil and extant taxa. At this moment, we did not have any reliable information on its ancestry and the degree of dwarfism, though the estimated body mass for this species implied that its body size seemed to be comparable to the continental extant muntjac species. In this paper, we present it as Muntiacini gen. et sp. indet. and refer to it as “the Ryukyu muntjac.” All of the specimens that were examined are stored at Okinawa Prefectural Museum and Art Museum (OPM), Naha, Japan.

*Sinomegaceros yabei* (femur N = 2, tibia N = 2): this fossil deer had a gigantic body size that was comparable to that of Irish elk (*Megaloceros giganteus*) [30, 43]. The skeletal remains were excavated from the Kumaishi-do Cave site (35°45″N, 137°′ 00″E, Fig. 1) in Hachiman-cho, Gujo city, Gifu Prefecture, central Japan. Most of the animal remains obtained from this site are isolated materials, but an incomplete skeleton that includes the skull has also been discovered [43]. The geological age of these fossils is the Late Pleistocene based on radiocarbon dating of the bones of cervids and proboscideans [44]. All of the specimens we sampled are stored at the Osaka Museum of Natural History (OMNH), Osaka, Japan.

Detailed information on the extant and fossil specimens is provided in Supplementary Information (SI) Tables S1–S3.

### 2. Thin sectioning and X-ray computed tomography (CT) scanning

We applied histological analyses of long bones to infer life history traits [8, 17-22].

We used 62 extant and 21 fossil long bones for histological analyses (Table 1). All specimens were then photographed and standard measurements were taken, following which thin sections of the midshaft of long bones were prepared based on the methodology described in previous studies [45, 46]. We produced 83 thin sections (62 sections for extant species, and 21 sections for extinct ones; see also detail in SI Table S1–S3). The thin sections were photographed using a digital film scanner (Pixus Mp 800, Canon) and analyzed using an Optiphot2-pol microscope (Nikon). Microscopic photographs were finally taken with a Nikon Df camera.

Neonatal specimens of sika deer (Honshu mainland: CBM-ZZ-757) and Reeves’s muntjac (CBM-ZZ-4974) were scanned using an experimental animal X-ray CT scanner (Latheta LCT-200, Aloka; 24-μm resolution, 80 kV, 0.2 mA) at Okayama University of Science, Okayama, Japan, to determine the initial diameter of the cortical bone and open medullary cavity in these species. Afterward, image segmentation and visualization were conducted using VG-Studio Max (Volume Graphics) version 3.1.

### 3. Age assessment

To allow for the comparison of growth patterns, we estimated ages as follows: for most of the extant deer samples, age at death was recorded in the museum inventory lists, which were assessed using tooth eruption or the number of dental cementum annuli in the tooth root (SI Tables S3 and S4). However, with fossil deer samples, long bones were unassociated with skulls and mandibles. Therefore, it was required to estimate the age at death from the number of LAGs found in the long bones. Several previous studies have reported a good correlation between the number of LAGs and the age at death in several vertebrate groups [47], while other studies indicated the limitation of LAGs as reliable ages due to loss of LAGs by bone remodeling and/or the expansion of the medullary cavity [e.g. 25, 48]. A study examined this correlation in extant deer species [red deer (*Cervus elaphus*)], which reported that although the number of LAGs in the tibia corresponded to the actual age, the number in the femur corresponded to the age before the deposition of the EFS [49]. Therefore, the relationship was examined first using extant deer between the actual age at death and the number of LAGs in bones (see SI for details). Results showed that the estimated age of the extant deer based on the number of LAGs, including those that appeared in EFS in both the femur and tibia was highly correlated with the actual age determined using the tooth cementum annuli (SI Figure S6 and Table S4). Furthermore, while some old adults who were over 8 years (HOUMVC-00037, CBM-ZZ-412, and URB-MAM-193) had fewer LAGs in their long bones than their actual age, diameters of their expanded medullary cavities were identical or even larger than bone diameters observed in neonates and fawns (<1 year old), proposing that the first (and also second in URB-MAM-193) LAG was eliminated in old individuals (SI Table S4). Thus, since the number of LAGs matched the actual age in extant sika deer, by counting the number of LAGs in their long bones, the age of individuals of the fossil deer was also estimated (SI Table S5). Secondary bone remodeling, which had erased any evidence of the primary bone tissue including LAGs, were seen only at the inner medullary surface and part of the cortex where ligaments were strongly attached (e.g., the labium laterale in the femur) in adults, indicating a good preservation of the primary bone tissues in most parts of the cortex.

### 4. Body mass estimation

The body mass during ontogeny was estimated from bone diameters at the LAGs [50]. Body mass estimation formulas for both the femora and tibiae were obtained and the formula that uses the mediolateral diameter (MLD) of the femora and tibiae was selected for use in this study (see SI for details). In addition to diameters at the LAGs, external diaphysial measurements [anteroposterior diameter (APD) and MLD] were collected and used to estimate body mass at death. Raw measurement data of diameters at the LAGs, with the estimated body mass used for growth curve modeling, are presented in Supplementary Table S13 (separately provided in excel format).

### 5. Comparison of growth patterns

Growth curves of body mass were constructed for each specimen that was older than yearling (i.e., those that had more than one LAG), which had at least four data points, including starting and terminal points. The exception to this was the extant Reeves’s muntjac, for which a fawn and yearling were included in the growth curve fitting because this species attained maturity early in life [35]. Before fitting the growth curves, neonatal body mass data were obtained for both the extant and fossil taxa as the starting point (SI Table S6 and Figure S7).

To fit growth curves to the obtained data, the number of LAGs was first transformed into the age in years. For each specimen, the body mass (*W*, in kg) was then formulated with age (*x*, in years) using the Gompertz curve, which is described below:

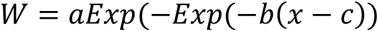

where *a, b*, and *c* represent the asymptote (i.e., the body mass when body growth ceases), growth rate, and inflection point, respectively. The Gompertz curve fitting was rationalized as this model showed the highest value of goodness of fit represented by the lowest AICc among the fitted growth curves (SI Table S7). Furthermore, the growth rate (*b*) and inflection point (*c*) were statistically compared among species/populations. First of all, the normality of the parameter distribution was confirmed by the Shapio-Wilk tests (for each species/population, *P* > 0.05). We then compared the growth rate and inflection point between the femur and tibia using extant deer samples by paired *t*-test. We found that the null hypothesis of no difference in these parameters between the femur and tibia was not rejected (for the growth rate, N = 15, *t* = 0.10, d.f. = 14, *P* = 0.92; for the inflection point, N = 15, *t* = 0.60, d.f. = 14, *P* = 0.56). Therefore, to increase the sample size we combined data from the femur and tibia for fossil samples, though we admit that further data are needed to clarify the difference in growth curves between the femur and tibia. We conducted one-way ANOVA and subsequent multiple comparisons using the Tukey-Kramer method to examine differences in the growth parameters among the eight species/populations, averaging the growth curve parameters from the femur and tibia for the extant species to avoid repeated sampling from the same individual.

Additionally, two-way ANOVA using bone type (i.e., femur or tibia) and species/populations as the factors was conducted. Growth curve parameters were averaged for each population/species to produce a representative growth curve for each group. All statistical analyses were conducted in JMP Pro 16.0 (SAS Institute Inc.).

### 6. Qualitative assessment of bone histology and external fundamental system (EFS) observation

To corroborate findings derived from estimated growth curves, we additionally investigated bone histology in detail. We observed the same histological sections used for the growth curve modeling. The nomenclature and definitions of bone tissues were based on the study by Francillon-Vieillot *et al*. [51] and Castanet *et al*. [52]. We focused on the following bone tissue characteristics: 1) the type of primary periosteal bone tissue and vascularization pattern as an indicator of the relative bone growth rate and 2) the presence of an external fundamental system (EFS). A primary periosteal bone tissue known as a fibro-lamellar (FBL) bone, which has a high level of laminar to plexiform vascularization (PV), is an indicator of fast-growth, whereas a parallel-fibered (PF) bone tissue is formed during periods of slower growth [e.g. 31]. Growth rate correlates with intensity and vascularization patterns [16, 53, 54], where lower levels of vascularization associated with the longitudinal orientation of canals indicated a slower bone depositional rate. Additionally, EFS is characterized by the PF bone tissue constituting without vascularization, thereby forming an edge of slow-growing bone tissues as a sign of somatic maturity [e.g. 48, 55, 56]. By counting LAGs within EFS, we estimated the ages at which EFS formation started (see SI for details). Somatic maturity was additionally investigated by the status of fusion of both proximal and distal epiphyses. Lastly, we conducted correlation tests between the age at which EFS development started and the inflection point (*c*), which marks the transition from the initial phase of exponential growth to the subsequent phase of asymptotic growth and is known to be correlated with the timing of sexual maturity in laboratory-reared mice [57]. For this purpose, we used the mid-values of EFS developmental age for each specimen. For example, OPM-HAN06-588 (*Cervus astylodon*) had the range of 9 – 10 years, therefore we used the value of 9.5 years as the age at which EFS development started. As for the extant specimens, we averaged the inflection point obtained from the femur and tibia growth curves. Non-parametric Spearman’s rank correlation coefficient (ρ) was calculated in three datasets (extant only, fossil only, and all individuals combined).

### 7. Comparison of survivorship curves

Since a change in mortality patterns also represents life history changes, we estimated survivorship curves of both extinct and extant deer populations. We obtained age data for the three extant sika deer populations that were used in the histological investigations either from published papers or through investigation of the cementum annuli of the incisor roots: Hokkaido mainland (N = 1,060) [58], Honshu mainland (N = 594) [59], and Yakushima Island (N = 74, cementum annuli of museum specimens). Histological analysis of cementum layers of the lower first incisors was conducted on Yakushima Island deer specimens housed in the University Museum of the University of Tokyo, the Tochigi Prefectural Museum, and the Hokkaido University Museum to identify their age at death. This analysis was done by a specialized laboratory (Matson Laboratory, Montana, U.S.A.) and we received age data with high reliability. The age data from three populations were based on randomly culled individuals and so were considered to be indicative of the age structure of actual extant populations. Because the hunting of female deer had been legally prohibited until 1994 and these deer populations were increasing in number at the time of culling (approximately from the late 1990s to early 2000s), the deer populations were less affected by the hunting pressure [59]. Additionally, we obtained mortality data of sika deer from Kinkazan Island from the published literature (N = 264) [60]. The Kinkazan population is protected for religious reasons and thus free from hunting pressure, which was also the case for *C. astylodon* and the Ryukyu muntjac inhabiting the Pleistocene Okinawa Island without human and other predators [60, 61]. The age data for extant Reeves’s muntjac from England was obtained from a published reference (N = 84)[62]. The age data of two fossil insular deer from Okinawa Island were also obtained from published references (*C. astylodon*: N = 45 [60]; the Ryukyu muntjac: N = 65 [63]).

Kubo et al. [60] estimated the age at death for *C. astylodon* based on the height of the lower third molar by applying the molar wear rate of extant sika deer. Variability of age estimation based on the molar wear rate was tested and discussed in previous studies [59, 60, 64], after which we adopted the most appropriate age estimates. Ozaki [63] estimated the age at death of the Ryukyu muntjac by applying the molar wear scoring method developed by Chapman *et al*. [62] for extant Reeves’s muntjac. Fossil *S. yabei* and extant Kerama Islands deer were excluded from these survivorship curve comparisons due to a lack of reliable age data.

To depict the survivorship curves of three culled sika deer populations, we constructed life tables following the method described in a reference [65], assuming stationary age distributions, and applied probit regression to smooth the age distributions. The age data for the other populations (Kinkazan sika deer, Reeves’s muntjac, and two fossil insular deer) were based on natural deaths, therefore the age class frequency was divided by the sample size to obtain a multiple of the frequency of mortality (dx). Life tables were then derived from this mortality frequency [65]. The survivorship curves were presented with an initial value of 1,000 individuals. The observed age class frequencies and derived life tables for each deer population are shown in SI Tables S11 and S12.

For more details on the materials and methods used in this study, see the SI and Supplementary Figures and Tables therein.

## Results

### Growth curve models and growth rate comparison using skeletochronology

The LAGs in the long bones of both extant and fossil deer were well preserved (SI Figure S1–4), allowing us to measure bone diameters at the LAGs. A growth curve of each species/population is presented in Figure 2A and SI Figure S10. Among the sika deer populations, deer from the two mainland areas (Honshu and Hokkaido mainland) showed faster growth than those from the two insular populations (Yakushima and Kerama Islands). Growth trajectories of body masses also corresponded with those obtained from investigations of culled individuals [66, 67] and through long-term field observations of wild deer (Agetsuma and Agetsuma-Yanagihara, personal communication), thereby validating our estimation method of body growth from skeletochronology. The extant Reeves’s muntjac showed the fastest growth, which was reported as well in the analysis of culled individuals that showed early sexual maturity at ca. 6 months [35].

**FIGURE 2.**
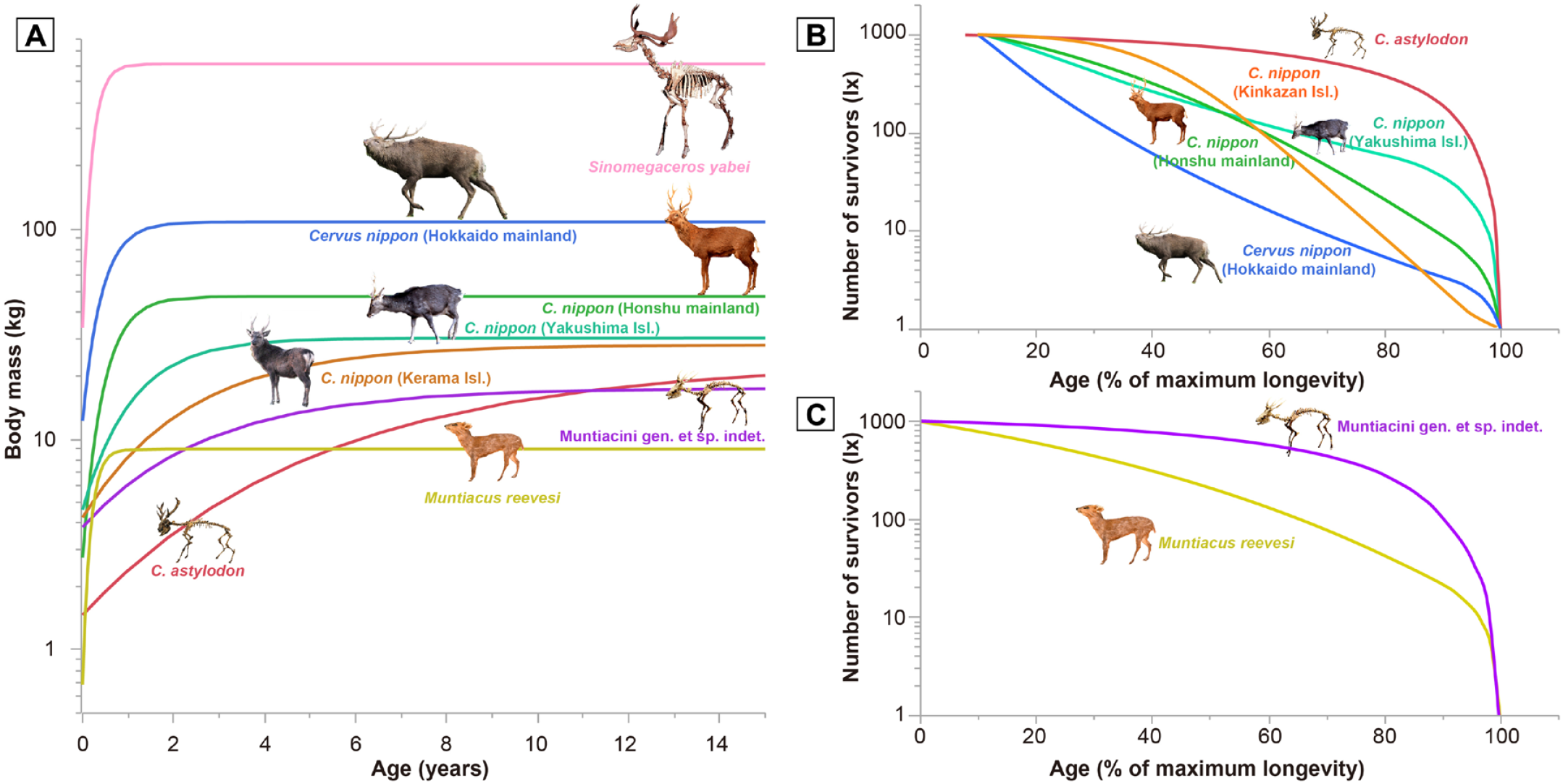
Growth (A) and survivorship curves (B-C) for studied Japanese deer. The insular deer have a significantly slower growth rate and a delayed growth plateau (somatic maturity) compared with the mainland deer (A), and an ecological shift exists in the mortality pattern between the insular and mainland deer (B, C). As the age estimation of *C. astylodon* was based on the lower third molar height, we confined the survivorship curves to juvenile and onward for *C. astylodon* and the sika deer (B).

Compared to the extant references, two fossil insular deer species (*C. astylodon* and the Ryukyu muntjac) showed remarkably slow and prolonged growth, whereas the Pleistocene giant deer (*S. yabei*), had fast growth, which suggested a growth pattern similar to other extant large deer species, such as moose [68].

Statistical comparisons of growth parameters revealed that both fossil and extant insular deer had significantly slower growth rates than Reeves’s muntjac and *S. yabei* (both *P* < 0.05, SI Tables S9, S10). However, growth rates of the four extant sika deer populations and fossil insular deer were not significantly different from each other (*P* > 0.05, SI Tables S9, S10), due to the transitional position of extant insular deer, although the limited numbers of possible samples may also play a role.

### Variation in bone tissue among deer populations

Histological characteristics of long bones corroborated the above statistical comparisons of growth curves. Figure 3 summarizes the main histological features of examined deer (also see SI Tables S1–S3).

**FIGURE 3.**
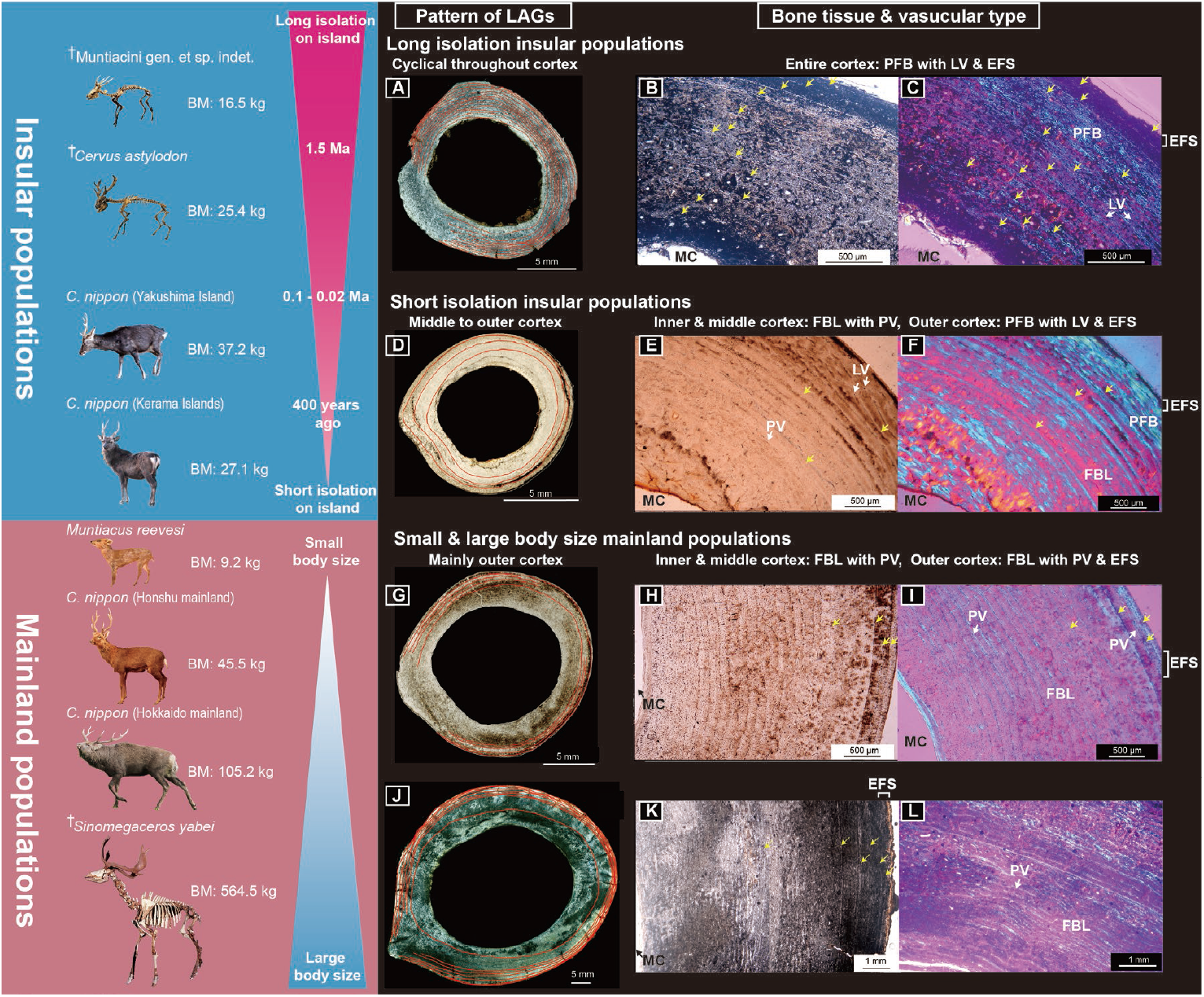
Bone histological features of studied Japanese deer. (A–C) Midshaft cross-section (A), the cortex under normal light (B), and the cortex under polarized light (C) of a femur of fossil *Cervus astylodon* (OPM-HAN07-1603). (D–F) Midshaft cross sections (D), the cortex under normal light (E), and the cortex under polarized light (F) of a femur of extant sika deer (*Cervus nippon*) from Yakushima Island (TPM-M-313). (G–I) Midshaft cross section (G), the cortex under normal light (H), and the cortex under polarized light (I) of a femur of extant sika deer from the Honshu mainland (CBM-ZZ-412). (J–L) Midshaft cross section (J), the cortex under normal light (K), and the cortex under polarized light (L) of a femur of fossil *Sinomegaceros yabei* (OMNH-QV-4062). Red lines in A, D, G, and J, and yellow arrows in B, C, E, F, H, I, and K indicate lines of arrested growth. The outer cortex is at the upper right of B, C, E, and F and on the right in H, I, and K. BM, body mass; EFS, external fundamental system; FBL, fibro-lamellar bone tissue; LAG, line of arrested growth; LV, longitudinal vascular canal; MC, medullary cavity; PFB, parallel-fibered bone tissue; PV, plexiform to laminar vascular canal. ^✝^Fossil cervids. Note: only fossil deer from Okinawa Islands had a distinctly longer-term geographical isolation history and exhibited bone tissue, indicating a slower growth rate [31].

As observed, all extant deer populations (i.e., the four extant sika deer populations, the extant Reeves’s muntjac), including the gigantic fossil *S. yabei*, exhibited a similar bone tissue structure to continental cervids [18, 48, 49], consisting of a fast-growing FBL bone with PV in most parts of the cortex (Figure 3G–L; SI Figures S2A–I, S3G– I, and S4A–O). Additionally, slow-growing PF bone tissue was observed in the outer cortex. This PF bone tissue was more extensively and notably developed in the two insular populations (sika deer from Yakushima and Kerama Islands), with LAGs at the inner to the middle cortex being tighter than in the mainland deer (Figure 3D–F, SI Figures S2D–F, and S3G–I, J–L), suggesting a slower growth rate among insular populations. Furthermore, the cortical bone tissues of two fossil insular deer (*C. astylodon* and Ryukyu muntjac) were different from those of extant deer and gigantic fossil deer (*S. yabei*), in that they were primarily PF bone tissue with multiple LAGs, thereby implying a much slower growth rate than other deer populations. Their bone tissues were also similar to those of the extinct dwarf bovid (*Myotragus balearicus*) and typical extant reptiles [e.g. 69]. Moreover, although LAGs were spaced evenly throughout the cortex (Figure 3A–C; SI Figures S1A–C and S3A–F), they became closer as they approached the periosteal surface in older individuals, which indicated a decrease in growth rate with age. Only a few areas of the innermost cortex showed a prevalence of the FBL bone tissue with primary osteons, which alternated with nonvascular PF bone tissue with LAGs, suggesting that the bone was deposited slowly during most periods of the lifetime. The slow growth rate was further supported by the observed vascularization pattern. Our fossil insular deer samples showed less vascularization and their vascular canals oriented longitudinally, which is not observed in other cervids and large mammals [18, 22]. In summary, the bone histological features of the fossil insular deer supported their slow growth rates.

Results also showed that all extant specimens with EFS had fused epiphyses both in the femora and tibiae (SI Figure S5, Table S3). As observed, the proximal epiphysis of the tibiae and femora fused later than the distal epiphysis. Moreover, EFS developed contemporaneously with proximal fusions. In two mainland sika deer populations, the timing of EFS development and proximal epiphyseal fusion showed a good correspondence with the age when the rate of growth decelerated (SI Figures S8, S9, Table S3), including the reported age of somatic maturity in the analyses of culled individuals [66, 67]. Notably, the extant sika deer from Kerama Islands also showed delayed EFS development and epiphyseal fusions compared with other investigated deer populations. Based on the observation of extant specimens, we further investigated the relationship between EFS development and epiphyseal fusion in fossil specimens (SI Figure S5, Tables S1, 2). Though most fossil specimens did not preserve both proximal and distal epiphyses, all specimens with fused proximal epiphysis had EFS and vice versa. Therefore, the correspondence of epiphyseal fusion and EFS was also held in fossil species as well. Findings revealed that two insular extinct deer showed much delayed EFS development (Min. 9 years, Max. 16 years for *C. astylodon*; Min. 5 years, Max. 11 years for the Ryukyu muntjac, see SI Tables S1, S2) than other investigated deer populations, suggesting a delay of somatic maturity.

Histological observation of delayed somatic maturity was further corroborated by the growth curve parameter. We found a significant positive correlation between the inflection point and the age at which EFS development started (extant only, ρ = 0.87, *P* = 0.0012; fossil only, ρ = 0.84, *P* = 0.0003; all data combined, ρ = 0.91, *P* < 0.0001; SI Figure S11). Though the inflection point was younger than the actual age when the body growth deaccelerated and the age at which EFS development started, the significant positive correlation indicated that the higher values of the inflection point can be used as an indicator of delayed somatic maturity. Statistical comparison of inflection points among studied deer showed that the inflection point was significantly greater in *C. astylodon* than in other deer populations (*P* < 0.05, SI Tables S9, S10), implying again that sexual maturity can be delayed in this species.

### Survivorship curve comparison among extant and fossil deer

To clarify life history differences in mortality patterns, we compared the survivorship curves of fossil *C. astylodon* and four other extant sika deer populations from the Hokkaido mainland, Honshu mainland, Yakushima Island, and Kinkazan Island (Figure 2B). Since the age estimation of *C. astylodon* was based on the lower third molar height, we confined survivorship curves to juvenile and onward for *C. astylodon* and the sika deer. A low mortality from the juvenile through to the prime age period characterized the survivorship curve of *C. astylodon*, followed by an increase in mortality during senescence, which appeared like an inversed L, and was categorized as a type I survivorship curve [70], including a characteristic of slow life species. At the other extreme, extant sika deer from the Hokkaido mainland showed relatively high mortality in the younger age class, which was followed by a more gradual decrease in the number of survivors from the prime age to senescence, thereby serving as a characteristic of type II survivorship curves. Results also showed that survival curves of sika deer from the Honshu mainland and Yakushima Island were located between those of *C. astylodon* and the Hokkaido sika deer, with a steady decrease in survivorship until the prime age, though the Honshu population showed an increased mortality rate after the prime age. Also, although the survivorship curve of the Kinkazan Island deer was clustered with Honshu and Yakushima Island populations, a lower mortality rate in the initial phase and a higher rate in the later phase was shown. This slight difference in the pattern is proposed to be associated with the lack of hunting pressure in the Kinkazan population. However, the intensity of hunting in the sika deer does not strongly affect overall survival curves as described in the Methods section. Furthermore, though the Kinkazan deer inhabited the predator-free island, their mortality pattern was different from that of the *C. astylodon*, implying a further advanced insular effect in the latter. A previous study reported that the maximum observed age of *C. astylodon* was 25 years [60], which was greater than that of the extant sika deer (17, 19, 17, and 21 years for the Honshu, Hokkaido, Yakushima Island, and Kinkazan Island populations, respectively; SI Tables S11, 12). We additionally compared survivorship curves between the Ryukyu muntjac and the extant Reeves’s muntjac (Figure 2C). As observed, the extant muntjac had a type II survivorship curve as in the extant sika deer from the Honshu mainland. Similarly, the Ryukyu muntjac had a type I survivorship curve, indicating that its mortality rate did not also increase until senescence, thereby implying a strong insular effect on life history traits in Okinawa Island again.

## Discussion

All interdisciplinary analysis results from growth rate comparison, histological investigation, and analysis of survivorship curves of both extant and extinct deer species supported our expectations that the evolution of a new combination of life history traits amounts to repeated establishments of a “slow life” in insular environments. Though the sample size of deer for each population was not very large, our findings are among the first to clarify the gradual shift to slow life histories for insular forms, placing the two fossil insular deer populations from Okinawa Island at the slow end of the fast-slow continuum.

It is possible to infer ecological processes of life history change from the current dataset obtained from deer populations. As observed in mainland deer populations, any modification of life history traits happened in mainland populations (Hokkaido and Honshu mainland sika deer, Reeves’s muntjac, and *S. yabei*), similar to the condition of continental relatives [18]. The Japanese Archipelago is considered “an island” in literature [e.g. 2]. Nevertheless, from the current dataset, the Japanese mainland deer is proposed not to be under a strong insular effect on life history traits. However, further investigation of sika deer populations from the Asian continent will shed light on the degree of insularity for the Japanese mainland sika deer.

We hypothesize a process of life history evolution on Japanese island deer as follows: the first step can be exemplified by sika deer on Kerama Islands. As observed, bone histological features showing signs of slow growth accompanied the smallest body size and slowest growth rate among understudied extant deer populations. These deer were introduced to Kerama Islands from the Kyushu mainland ca. 400 years ago by humans (Okinawa Prefectural Board of Education 1996). Fast changes in body size in less than 6,000 years have also been documented previously in red deer from Jersey [71]. Furthermore, domestic cattle that were introduced to Amsterdam Island experienced a rapid shrinking of body size in ca. 110 years [72, 73]. Though changes in growth trajectories might exist, these changes were not revealed in these interesting populations. With the Kerama deer, this drastic change in growth trajectory is proposed to have occurred within 400 years, which equates to ca. 80 generations (assuming a generation time of 5 years for the sika deer). The change in the growth trajectory of the Kerama deer is therefore proposed to have been brought about by phenotypic plasticity rather than a genetic modification in response to the less abundant understory vegetation in the subtropical evergreen broadleaved forest, accounting for the poor nutritional status of the Kerama deer [74], which was supported by the observation of a possible malnourished individual (URB-MAM-55; see SI for details). Lister [26] originally hypothesized the importance of phenotypic plasticity in the initial stage of insular dwarfism, and behind this would be epigenetic mechanisms controlling body growth, which respond to habitat environments [75, 76]. These mechanisms should be a challenging theme in the study of insular evolution. Another topic to be clarified in the Kerama deer would be the life history shift observed in its demographic aspect. It is therefore relevant to obtain reliable mortality data for the Kerama deer for this purpose.

The Yakushima Island deer illustrates the next step in life history evolution, which also showed a slower growth rate and intermediate histological features, coupled with a moderate shift in survivorship curves toward those of slow life species. Contrary to the situation of the Kerama deer, the slower growth rate of Yakushima deer is proposed to have a genetic basis because; 1) Yakushima deer is not malnourished and have enjoyed a recent population increase and 2) some of their macromorphological features have a genetic basis [36, 38]. Long-term field observations of the Yakushima deer also supported our finding that life history traits of this population were shifting to slow life (Agetsuma and Agetsuma-Yanagihara, personal communication).

The clearest evolutionary transformation was shown in the two fossil deer species isolated from Okinawa Island over 1.5 Ma. Their growth patterns, bone histological characteristics, and mortality patterns were different from other deer populations. As with the Kerama deer, it was proposed that phenotypic plasticity partially influenced their growth trajectories. However, we do not currently have reliable sources of information on the nutritional status of the fossil deer, except for their dietary habits [61, 64]. Nevertheless, reports of healed bone fractures on several leg bones from a museum collection of fossil insular deer species on the Okinawa Island have been recorded, which implies that this deer survived long enough to recover from serious leg injuries. These findings refute the possibility that fossil Okinawa deer were malnourished and their body growth was suppressed due to a lack of resources. It was also notable that the same trend was observed for the fossil deer and muntjac, further supporting the shift toward slow life as an adaptation to the insular environment. This process is proposed to have started after their initial settlement on the Okinawa Island in part through phenotypic plasticity of growth trajectory due to a lack of sufficient foods during growing periods. A life history change would then have become genetically fixed through the natural selection of more slow life individuals, which occurred in the Yakushima Island deer. The outcome will have been the development of extremely slow life animals, despite their small body size. Unfortunately, this life history trait will have made these species vulnerable to human exploitation because animals having slow life histories have overall lower population recruitment than fast-life animals and their population recruitment is most declined by the increase of adult mortality rate [13, 70]. Consequently, the two Okinawa deer species became extinct at the time of or soon after the Paleolithic human arrival to the Okinawa Islands [77, 78], probably due to hunting. This extinction applies to other island dwarfs that became extinct during the Pleistocene era [79, 80, but see 81]. Recently, Rozzi et al. [82] analyzed body size changes and extinction risk of over 1,500 island mammals worldwide and clarified that insular dwarfs and giants are susceptible to extinction by the arrival of modern humans. It is plausible that behind the extinction there could be a life history shift toward the slow life in these insular extreme forms.

Our findings also agreed with worldwide observations of life history changes in insular mammals previously documented (Figure 4). A living cervid (*Odocoileus hemionus*) from Blakely Island showed slower growth rates than those from the mainland population [10]. As the tooth eruption was delayed in the Blakely Island deer, this life history change is proposed to have a genetic basis, similar to the case of the Yakushima Island deer. With longer isolation but variable island sizes, the life history response varied among mammals from Mediterranean islands. A fossil dwarf hippo (*Hippopotamus minor*) from Cyprus showed no evident modification in bone histology [19], though their life history traits remained unclarified. In contrast, a fossil cervid (*Candiacervus*) from Crete showed slight modifications toward a slower life history [18, 25]. The Sicilian pygmy elephant (*Elephas falconeri*) also showed a modification in life history. This species was observed to have a fast life history as the age distribution of its fossil assemblage was skewed to calves and immature individuals [83]. Nevertheless, a histological study suggested that this pygmy elephant had a much slower growth rate than its mainland relatives, and sexual maturity was within the range of extant elephants [84], thus this contradiction requires a further synthetic study on this species. The differences in the life history response among Mediterranean mammals, however, are unsurprising, given that these Mediterranean islands were much bigger than those for which we report life history changes (i.e., the Blakely Island, Kerama Islands, Yakushima Island, Okinawa Island, and Mallorca Island). The most explicit change in life history was observed in *Myotragus balearicus* from the Mallorca Island, which underwent an exceptionally long time of isolation (5.2 Ma) in predator-free environments. *Myotragus* showed a dramatic decrease in bone growth rate and an evolution toward a slow life history [8, 21, 23], implying the commonality of this evolutionary change with the Okinawa fossil deer. These results propose that both island size and duration of isolation, with smaller islands and longer isolation, strongly affected the degree of modification in life history, thereby resulting in life history changes toward the slow life.

**FIGURE 4.**
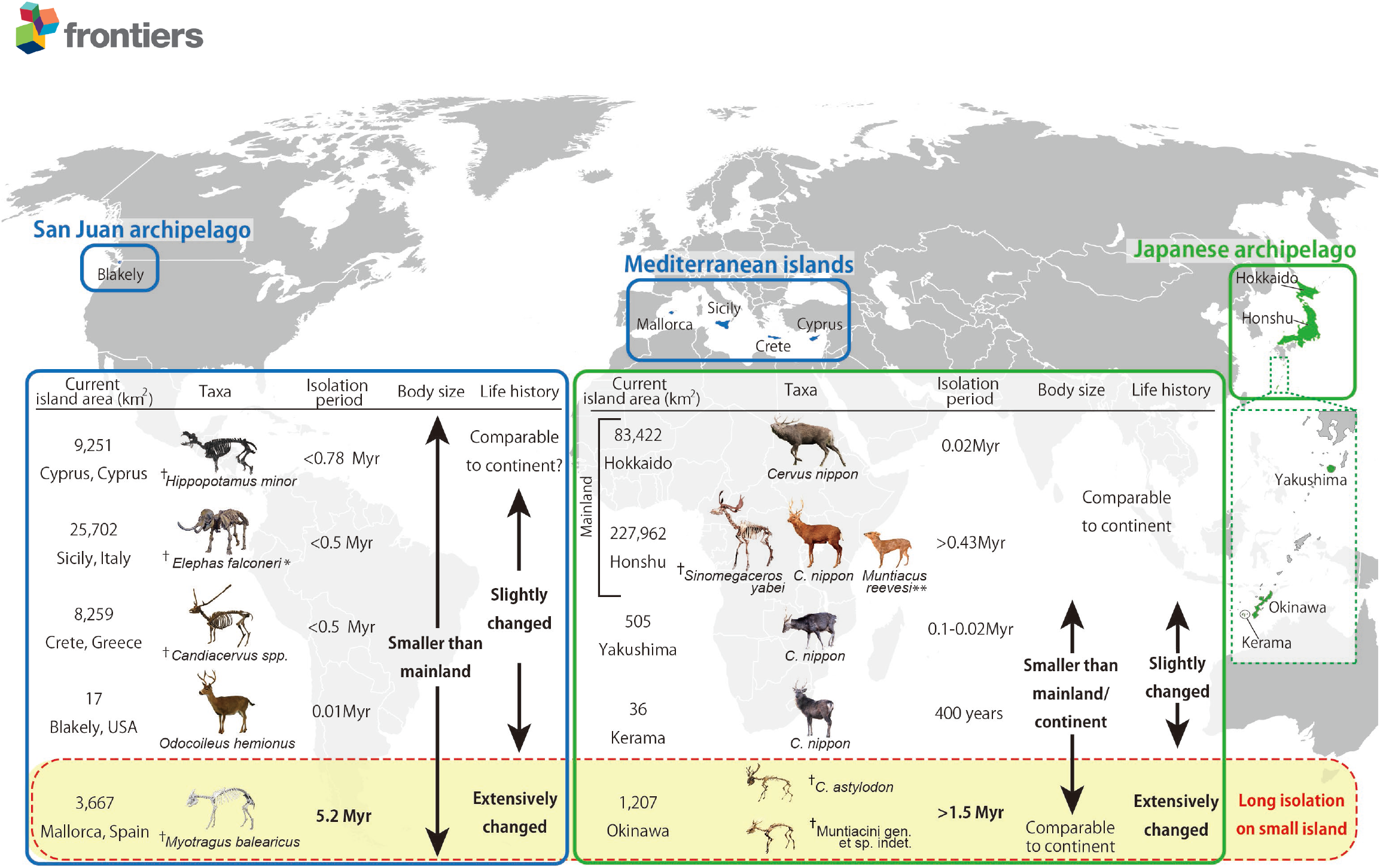
The Diagram summarizes evolutionary processes, the relationship between insularity, body size, and life history observed in extant and fossil large mammals. Cervid species/populations of the Japanese Archipelago were investigated in this study and information on animals from other islands we obtained from references: *Odocoileus hemionus* [10], *Hippopotamus minor* [2], *Candiacervus* spp. [2], *Elephas falconeri* [2, 85], *Myotragus balearicus* [8]. Photographs of fossil mammals in Mediterranean islands courtesy of Gunma Museum Natural History (*Elephas falconeri*), Alexandra van der Geer (*Hippopotamus minor* and *Candiacervus* spp.), and Meike Köhler (*Myotragus balearicus*). ^✝^Fossil taxa. ^*^ Life history of *Elephas falconeri* was disputed [83, 84]. ^**^ *Muntiacus reevesi* was an artificially introduced animal in the 1980s. Therefore, the isolation period of >0.43 Myr is inapplicable to it [35].

## Conclusions

The extant Japanese sika deer can modify their life history traits through a combination of phenotypic plasticity and natural selection processes. This ability underlies the process of insular dwarfism also recorded in fossil deer species (*C. astylodon* and the Ryukyu muntjac) from the Okinawa Island, which were estimated to have a slow life history. After combining results from the present and previous studies on the life histories of insular mammals, we conclude that both island size and duration of isolation strongly affect the degree of modification in life history traits, with longer isolation on smaller islands resulting in more prominent life history changes. We also demonstrated that the evolution of a gradual and transitional shift toward the slow life history in large mammals from the mainland to insular populations was the main life history change accompanying insular dwarfism, which makes the insular dwarfs vulnerable to human exploitation.

## Data Availability Statement

All data are available in the main text and electronic supplementary information.

## Ethic statement

Ethical review and approval were not required for this study used museum specimens and the investigations for the specimens were officially approved by the curators.

## Author contributions

SH, MOK, and MF designed the research. SH, HT, and TN, prepared bone histological sections, took measurements and created visualizations. MOK, MI, and TS collected ecological data. MOK conducted statistical analyses. SH, MOK, MSV, and MF drafted the original manuscript. All authors also contributed to reviews and edits of the manuscript and approved the final manuscript.

## Acknowledgments

We thank the following curators for access to museum collections under their care: Shimoinaba, S. (Natural History Museum and Institute, Chiba, Japan), Hayashi, T. (Tochigi Prefectural Museum), Eda, M. (The Hokkaido University Museum), Usami, K. (Okinawa Prefectural Museum and Art Museum). Oshiro I. is also acknowledged for his advice on the fossil Okinawa deer and support during this study. Furthermore, we thank Ochiai, K. and Shimoinaba, S. for providing ecological data on sika deer and Reeves’s muntjac in Honshu. Agetsuma, N. and Agetsuma-Yanagihara, Y. are appreciated for sharing data from their long-term field survey of the Yakushima deer. We are grateful to Nakamura, K. and Nomura, H. as well for technical assistance in thin-sectioning, and Kodaira, S. and Kodama, R. for CT scanning. We appreciate Kawada, S. (National Museum of Nature and Science, Tokyo), van der Geer, A. (Naturalis Biodiversity Center), Köhler, M. (ICP), Takakuwa, Y. (Gunma Museum of Natural History), Kikuchi, H. and Umemura, Y. for providing photos of *Odocoileus hemionus, Hippopotamus minor, Candiacervus* spp., *Myotragus balearicus, Sinomegaceros yabei, Elephas falconeri*, and sika deer (Yakushima Island and Hokkaido mainland, respectively). We are grateful for the comments of Weisbecker, V. and Köhler, M. on the earlier version of this manuscript.

