## Supplementary information for "Variation and process of life history evolution in insular dwarfism as revealed by a natural experiment"

**Methodological details**

**Validity of using the number of lines of arrested growth (LAGs) to estimate age**

We first checked the correlation between the numbers of LAGs observed in the long bones and the actual ages at death assessed by the numbers of dental cementum annuli formed in the tooth roots using 20 extant sika deer from both mainland and island localities (Table S4). Since the date or season of death had been recorded for all of these samples together with the age at death, we counted how many times deer experienced winter (i.e., the season when LAGs are usually formed) and compared this with the number of LAGs. The scatter plot clearly showed that, with the exception of four individuals, the number of LAGs equaled the number of winter seasons the deer experienced in their lifetime (Figure S6). The exceptions to this were one sika deer from Hokkaido mainland (HOUMVC00037), one sika deer from Honshu mainland (CBM-ZZ-412), and two sika deer from Kerama Islands (URB-MAM-55 and -193). HOUMVC00037, CBM-ZZ-412, and URB-MAM-193 had fewer LAGs (7, 7, and 8, respectively) than their actual age at death (8, 8, and 10, respectively). Based on the expansion of the medullary cavity in these old individuals, it seemed likely that the first (and also second in URB-MAM-193) LAG had been eliminated, and this was supported by the estimated body mass at the first LAG, which was greater than that of younger deer of the same population and sex (based on a comparison of HOUMVC00037 with HOUMVC00033, CBM-ZZ-412 with CBM-ZZ-218, and URB-MAM-193 with URB-MAM-211). Therefore, we considered that these three individuals had lost their initial growth information. In the case of URB-MAM-55, the number of LAGs (7) was greater than the actual age at death (4). This individual had limb bones without epiphyseal fusion, indicating that the bones were still actively growing in a longitudinal direction. However, observation of the long bone histology showed that appositional growth was decelerated and marked with many LAGs. Therefore, it is possible that this deer was malnourished and its growth was arrested intermittently even during the growing season. Consequently, we excluded this individual from the following growth curve analysis.

For the extant Reeves’s muntjac specimens, the age at death was estimated based on tooth eruption and the molar wear condition. Assessment of the correspondence between the number of LAGs and the age assessed by tooth eruption and wear for Reeves’s muntjac showed that the two were perfectly matched. Therefore, for this species, we considered the age assessed by tooth eruption and wear to be the actual age at death.

**Age assessment from LAGs**

We estimated the age at death of the fossil specimens from the number of LAGs. However, several specimens showed explicit expansion of the medullary cavity, which may have erased the record of early growth. Therefore, we also estimated the number of lost LAGs due to bone remodeling. Several estimation methods have been proposed in paleo-histological studies [1-3]. Here, we applied a simple method that uses a reference specimen with no apparent bone remodeling or expansion of the medullary cavity and compares this with specimens with possible dissipation of the initial LAGs. We chose this method because backward estimation from fitted growth curves [1] resulted in overestimation of the number of lost LAGs, generating a slower growth rate for the fossil taxa. Therefore, to yield a more conservative comparison with extant taxa, we estimated the number of lost LAGs through comparisons with younger specimens of the same taxa that were well preserved. Table S5 summarizes the estimation of lost LAGs based on comparisons with the reference specimens. The number of lost LAGs was small in all cases except for the femora of *C. astylodon*, which exhibited a more obvious expansion of the medullary cavity, resulting in up to seven LAGs potentially being lost (in OPM-HAN06-155).

Among the extant specimens we analyzed, only one sika deer (Kerama Islands, URB-MAM-34) did not have a reliable age based on the tooth cementum annuli. This specimen had eight LAGs in both the femur and tibia with completely fused epiphyses. Comparison of this specimen with other Kerama deer specimens indicated that one LAG had been erased by bone remodeling. Therefore, the age at death of this specimen was estimated to be 9 years old.

Raw data of LAG and external diaphysial measurements of specimens used in the histological analyses, together with estimated body mass is presented in Table S13 (Separate Excel file).

**Body mass estimation**

In addition to measuring the bone diameters at the LAGs, we measured the external diaphysial measurements [anteroposterior diameter (APD) and mediolateral diameter (MLD)] and used these to estimate the body mass at death. The body mass estimation formulae for both the femur and tibia were obtained from a reference [4]. In order to avoid extrapolation of data that beyond the original data range of regression model, we used the interspecific regression models to estimate the body mass through ontogeny, which can cover the wide range of body mass in the current dataset. It is also rationale by following reasons: 1) by adopting a single regression model to all specimens we can reduce model uncertainties emerging from applying different models to respective species, and 2) though we admitted that ontogenetic allometry (allometry observed in an individual) is different from static allometry (allometry observed in a group of full adult individuals) [5], we found our fitted growth models showed good correspondences with previously reported body mass changes in extant sika deer [6, 7] and Reeves’s muntjac [8]. The following linear regression equations of body mass against limb bone metrics for extant artiodactyls were used:

Log_10_ (body mass in kg) = 0.8737 + 2.7585*log_10_ (femur APD in cm)

Log_10_ (body mass in kg) = 0.8915 + 2.8771*log_10_ (femur MLD in cm)

Log_10_ (body mass in kg) = 0.9834 + 2.9356*log_10_ (tibia APD in cm)

Log_10_ (body mass in kg) = 0.8689 + 2.7672*log_10_ (tibia MLD in cm)

We compared the two formulae for each of the femur and tibia and selected those that produced the least discrepancy in estimated body mass with the femur and tibia metrics of extant cervids. This resulted in the formulae using MLD being selected for both bones.

Accurate fitting of a growth curve requires knowledge of the starting point (i.e., the neonatal body mass). For extant sika deer (Honshu and Hokkaido mainland, Yakushima Island) and Reeves’s muntjac, we collected neonatal or perinatal body mass data from the literature [9] or zoo reports. However, the neonatal body mass was not known for sika deer from Kerama Islands. Therefore, we used the value of sika deer from Yakushima Island for this population because of their similar body sizes as adults. For the fossil taxa, we estimated the neonatal body mass using the following adult–neonate body mass regression equation based on data from 121 extant artiodactyls in the PanTHERIA database [10; http://esapubs.org/archive/ecol/E090/184/] (Figure S7):

Log_10_ (neonatal body mass in g) = −0.3009 + 0.806*log_10_ (adult body mass in g)

The adult body masses of the fossil cervids, which were obtained from the external diaphysial measurements of the femora and tibiae, were averaged for each species and substituted into the regression equation to predict the neonatal body mass. The estimated neonatal body masses of the fossil cervids are presented in Table S6.

**Growth curve modeling**

For both the extant and fossil samples, all individuals that were older than yearlings were used for growth curve fitting, which had at least four data points in fitting growth curves. In addition, we included a fawn and a yearling in the growth curve fitting for extant Reeves’s muntjac. We tested the goodness of fit of the growth curves comparing four representative growth curves frequently used in growth analyses: a logistic curve with three parameters, a logistic curve with four parameters, a Gomperz curve with three parameters, and a Gomperz curve with four parameters. A growth model of von Bertalanffy was not used because for some individuals it produced unusual values of asymptotes or parameters did not converge. The test on the good ness of fit was done by “Fit Curve” platform provided by JMP (ver. 16), assigning the estimated body mass (either femur- or tibia-based estimate) being a response variable, the age in years as a regressor variable, and the individual specimen number as a grouping variable (i.e. curve-fitting was done individually for ontogenetic body mass series). We found that a Gompertz curve with three parameters had the lowest AICc values among the four models (Table S7). Therefore, the Gompertz curve with three parameters was fitted to the longitudinal body mass data for each individual using the formula:

$$W=aExp(-Exp(-b\left( x-c \right))$$

where *W* is the body mass in kg, *x* is the age in years, and the parameters *a*, *b*, and *c* represent the asymptote, growth rate, and inflection point, respectively. We then obtained growth curves for each species/population and each leg bone (femur or tibia) (Figures S8 and S9, Table S8). Our estimated growth curves based on the bone diameter measurements at LAGs were in good correspondence to published growth curves of the sika deer populations (Hokkaido and Honshu mainland populations) and the Reeves’s muntjac population, which were based on body mass of culled individuals [6-8]. Also, the growth pattern was comparable to that obtained by long-term field observations of Yakushima deer (Agetsuma and Agetsuma-Yanagihara, personal communication). Therefore, our estimation method of body growth from skeletochronology was validated.

**Supplementary Figures**

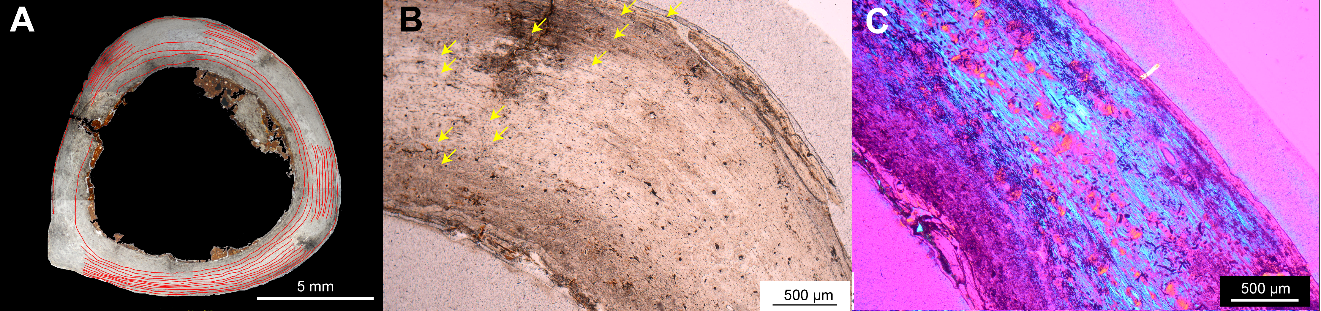

**Figure S1. Bone histologies of the femora of fossil Ryukyu muntjac. (A–C) Midshaft cross section in normal light (A), the cortex under normal light (B), and the cortex under polarized light (C) of femora of the Ryukyu muntjac (Muntiacini gen. et sp. indet.; A and B: OPM-HAN06-163; C: OPM-HAN07-1618). The red lines in A and the yellow arrows in B indicate lines of arrested growth (LAGs). The outer cortex is in the upper right in B and C. Note: LAGs appear at narrow intervals in the inner cortex in the Ryukyu muntjac, which also has an abundance of parallel-fibered bone tissue (blue tissues in C).**

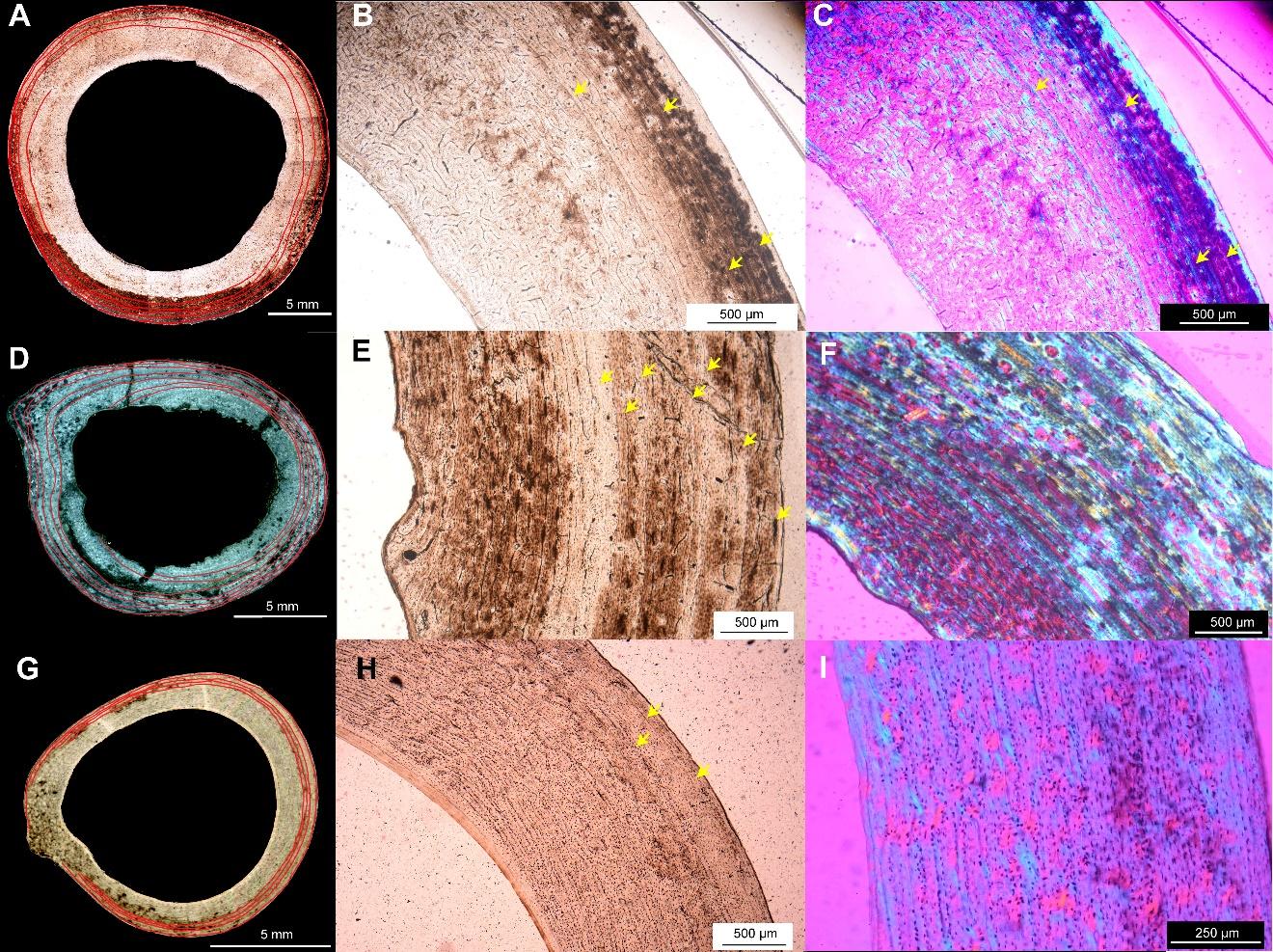

**Figure S2. Bone histologies of the femora of extant Japanese cervids. (A–C) Midshaft cross section in normal light (A), the cortex under normal light (B), and the cortex under polarized light (C) of a femur of sika deer (*Cervus nippon*) from Hokkaido mainland (HOUMVC00035). (D–F) Midshaft cross section under normal light (D), the cortex under normal light (E), and the cortex under polarized light (F) of a femur of sika deer from Kerama Islands (URB-MAM-183). (G–I) Midshaft cross section under normal light (G), the cortex under normal light (H), and the cortex under polarized light (I) of a femur of Reeves’s muntjac (*Muntiacus reevesi*) from Honshu mainland (CBM-ZZ-2646). The red lines in A, D, and G and the yellow arrows in B, E, and H indicate lines of arrested growth (LAGs). The outer cortex is on the right in E and I and in the upper right in B, C, F, and H.**

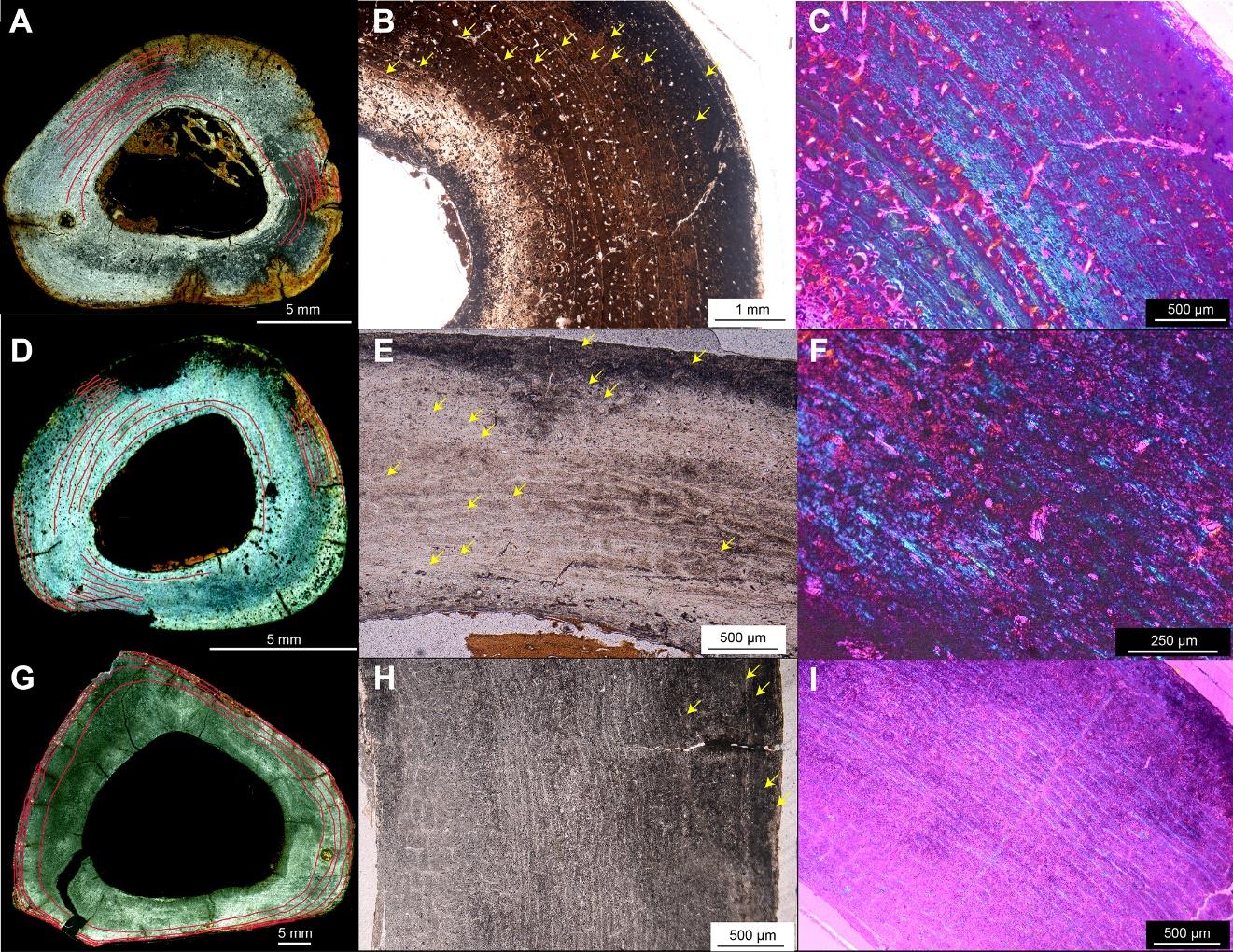

**Figure S3. Bone histologies of the tibiae of fossil Japanese cervids. (A–C) Midshaft cross section under normal light (A), the cortex under normal light (B), and magnified view of the cortex under polarized light (C) of a tibia of *Cervus astylodon* (OPM-HAN07-1585). (D–F) Midshaft cross section under normal light (D), the cortex under normal light (E), and magnified view of the cortex under polarized light (F) of a tibia of the Ryukyu muntjac (Muntiacini gen. et sp. indet.; OPM-HAN07-1599). (G–I) Midshaft cross section under normal light (G), the cortex under normal light (H), and the cortex under polarized light (I) of a tibia of *Sinomegaceros yabei* (OMNH-QV-4068). The outer cortex is in the upper right in B, C, F, and I, at the top in E, and on the right in H.**

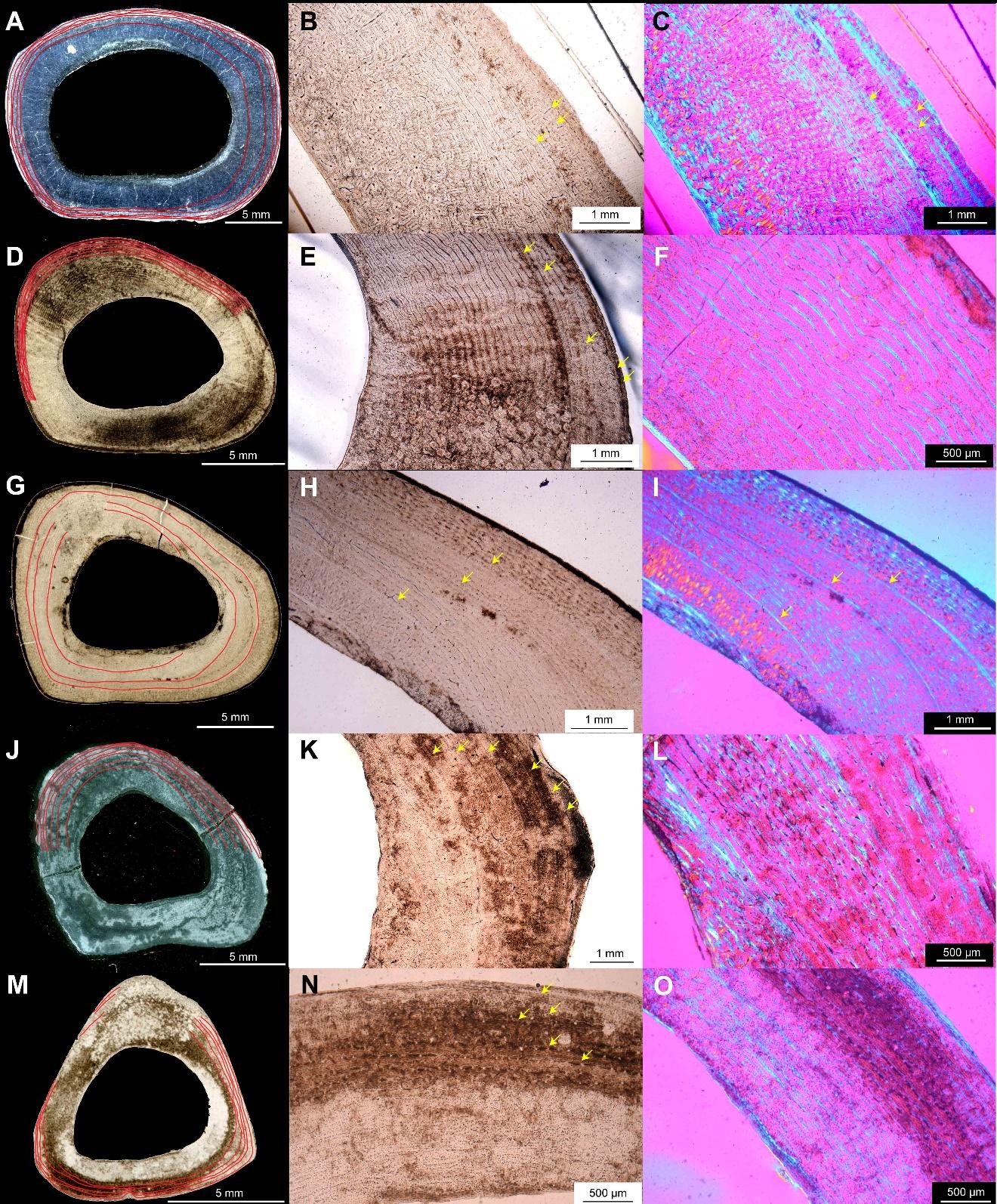

**Figure S4. Bone histologies of the tibiae of extant Japanese cervids. (A–C) Midshaft cross section under normal light (A), the cortex under normal light (B), and the cortex under polarized light (C) of a tibia of sika deer (*Cervus nippon*) from Hokkaido mainland (HOUMVC00035). (D–F) Midshaft cross section under normal light (D), the cortex under normal light (E), and the cortex under polarized light (F) of a tibia of sika deer from Honshu mainland (CBM-ZZ-412). (G–I) Midshaft cross section under normal light (G), the cortex under normal light (H), and the cortex under polarized light (I) of a tibia of sika deer from Yakushima Island (TPM-M-313). (J–L) Midshaft cross section under normal light (J), the cortex under normal light (K), and the cortex under polarized light (L) of a tibia of sika deer from Kerama Islands (URB-MAM-183). (M–O) Midshaft cross section under normal light (M), the cortex under normal light (N), and the cortex under polarized light (O) of a tibia of Reeves’s muntjac (*Muntiacus reevesi*) from Honshu mainland (CBM-ZZ-2646). The outer cortex is in the upper right in B, C, E, F, H, I, K, L, and O at the top in N.**

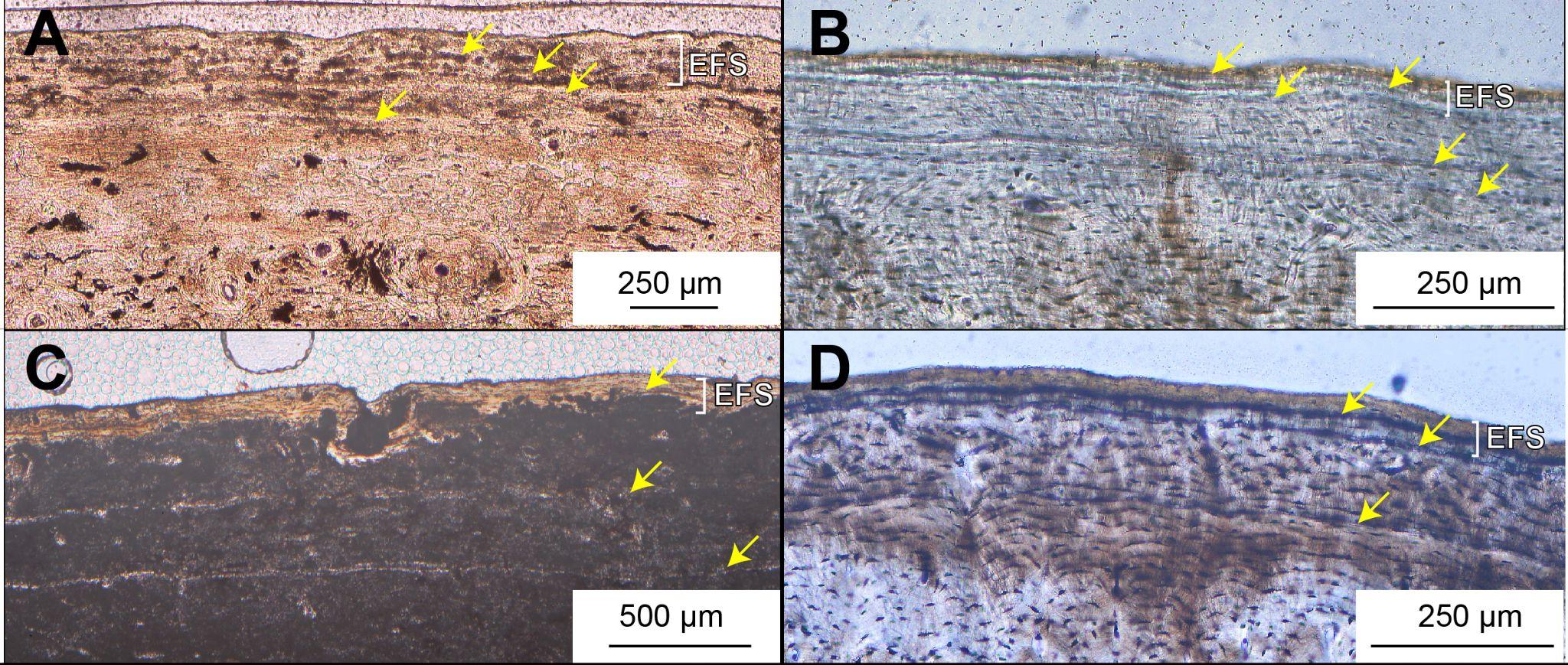
 **Figure S5. External fundamental system (EFS) of extinct and extant cervids. White parentheses indicate the presence of EFS. Yellow arrows show LAGs. (A) Cortex of a Ryukyu muntjac femur (Muntiacini gen. et sp. indet.; OPM-HAN07-1618). (B) Cortex of a sika deer femur from Yakushima Island (TPM-M-316). (C) Cortex of a *Sinomegaceros* *yabei* tibia (OMNH-QV-4068). (D) Cortex of a sika deer femur from Honshu mainland (CBM-ZZ-412).**

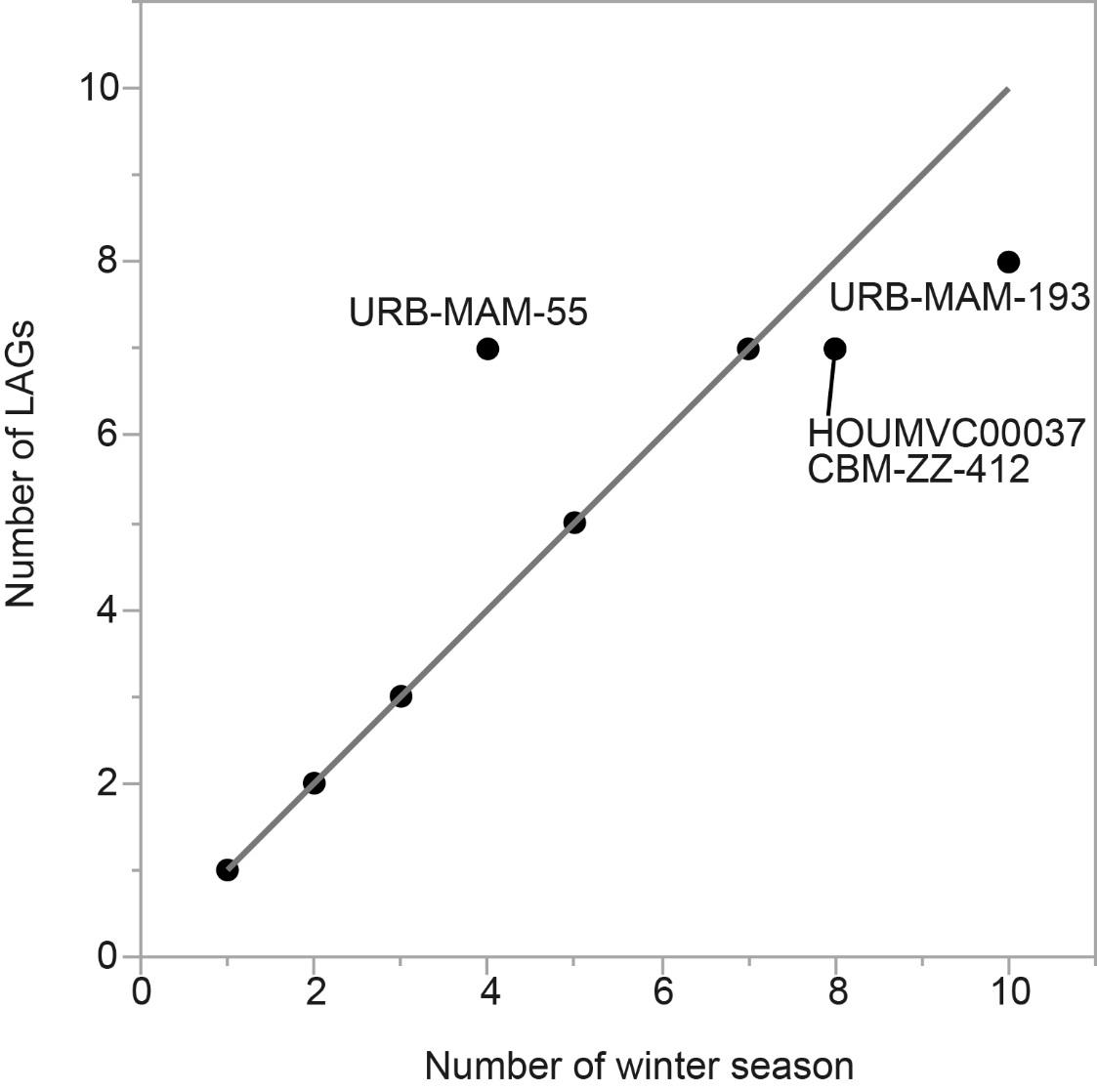

**Figure S6. Scatter plot showing the correlation between the number of winter seasons a deer experienced during its lifetime and the number of lines of arrested growth (LAGs). The raw data are presented in Table S5. There was a perfect match between the two variables in extant sika deer (*Cervus nippon*) of known age, with the exception of four individuals (HOUMVC00037, CBM-ZZ-412, URB-MAM-55, and URB-MAM-193). The solid line represents x = y.**

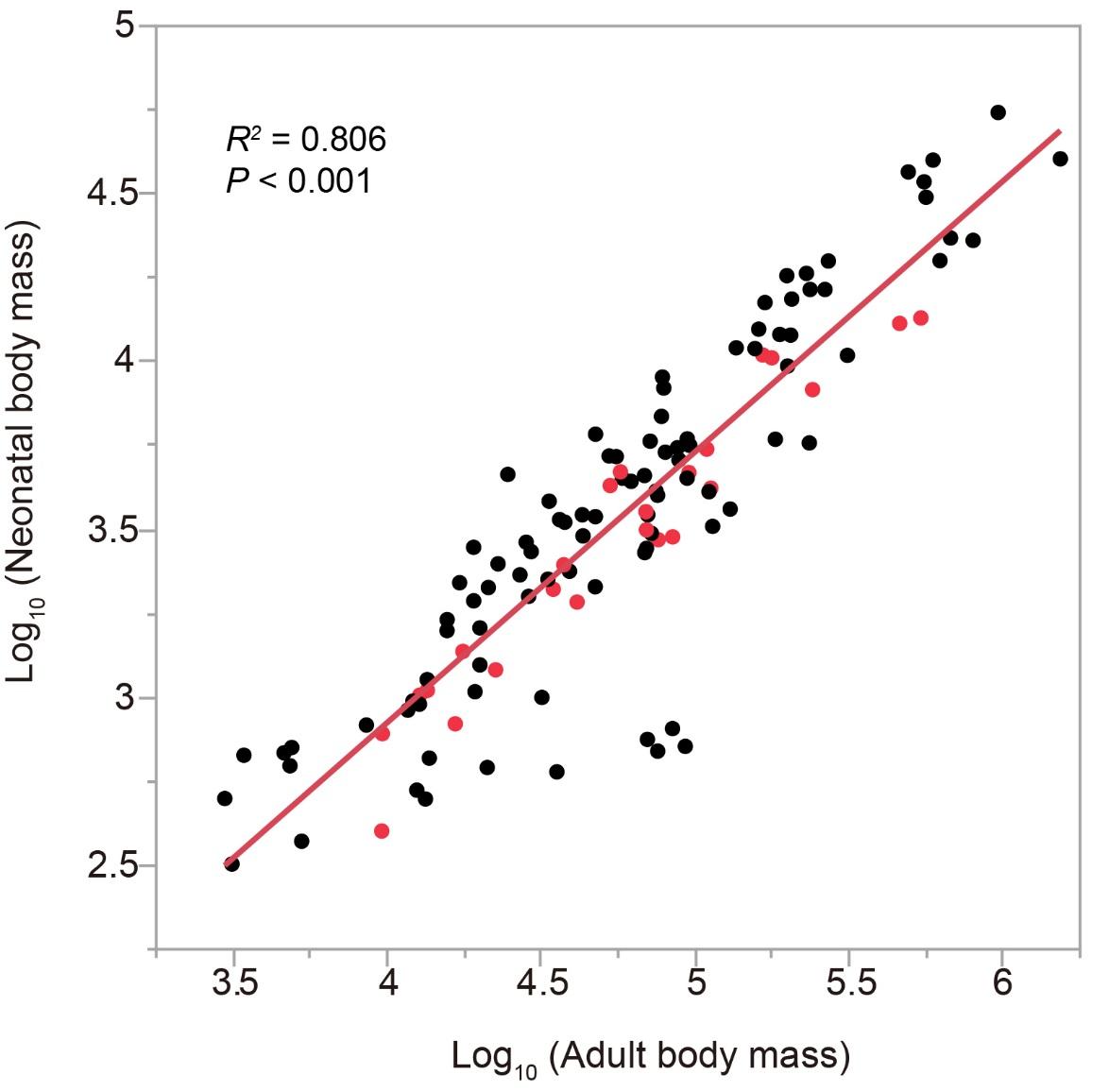

**Figure S7. Scatter plot showing the correlation between adult and neonatal body mass in extant artiodactyls (n = 121). The solid line represents the linear regression line. The red dots indicate Cervidae species. Data were obtained from PanTHERIA [10; http://esapubs.org/archive/ecol/E090/184/]. The regression equation was then used to estimate the neonatal body mass of three fossil cervids (see Table S6).**

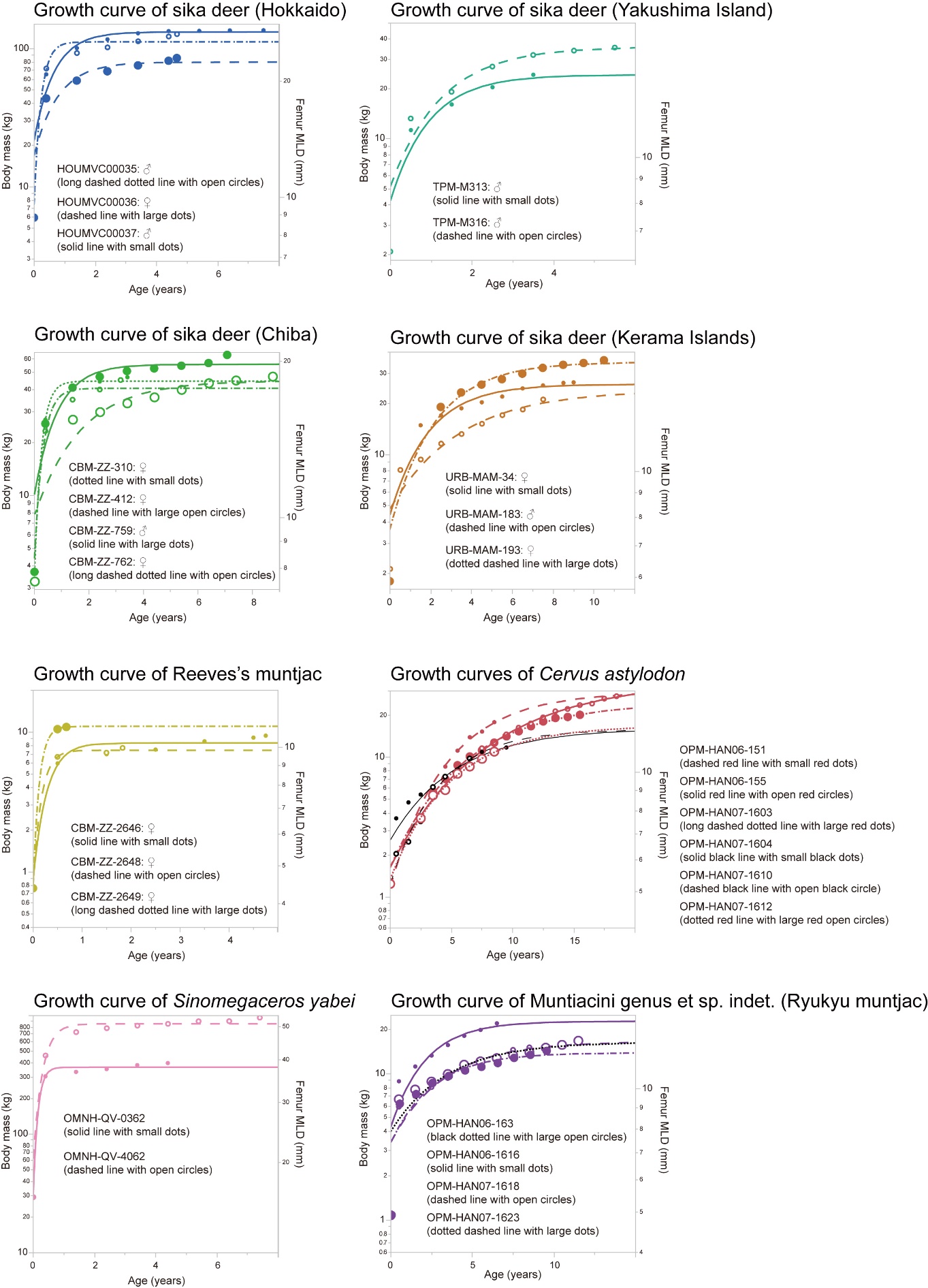
**Figure S8. Growth curves of body size estimates, i.e. the femur mediolateral diameter (MLD) in the right axis and body mass estimated from the femur MLD in left Y-axis, of eight cervid populations. The Gompertz curves are fitted and the parameters for which (*a*: asymptote; *b*: growth rate; and *c*: inflection point) are shown in Table S8.**

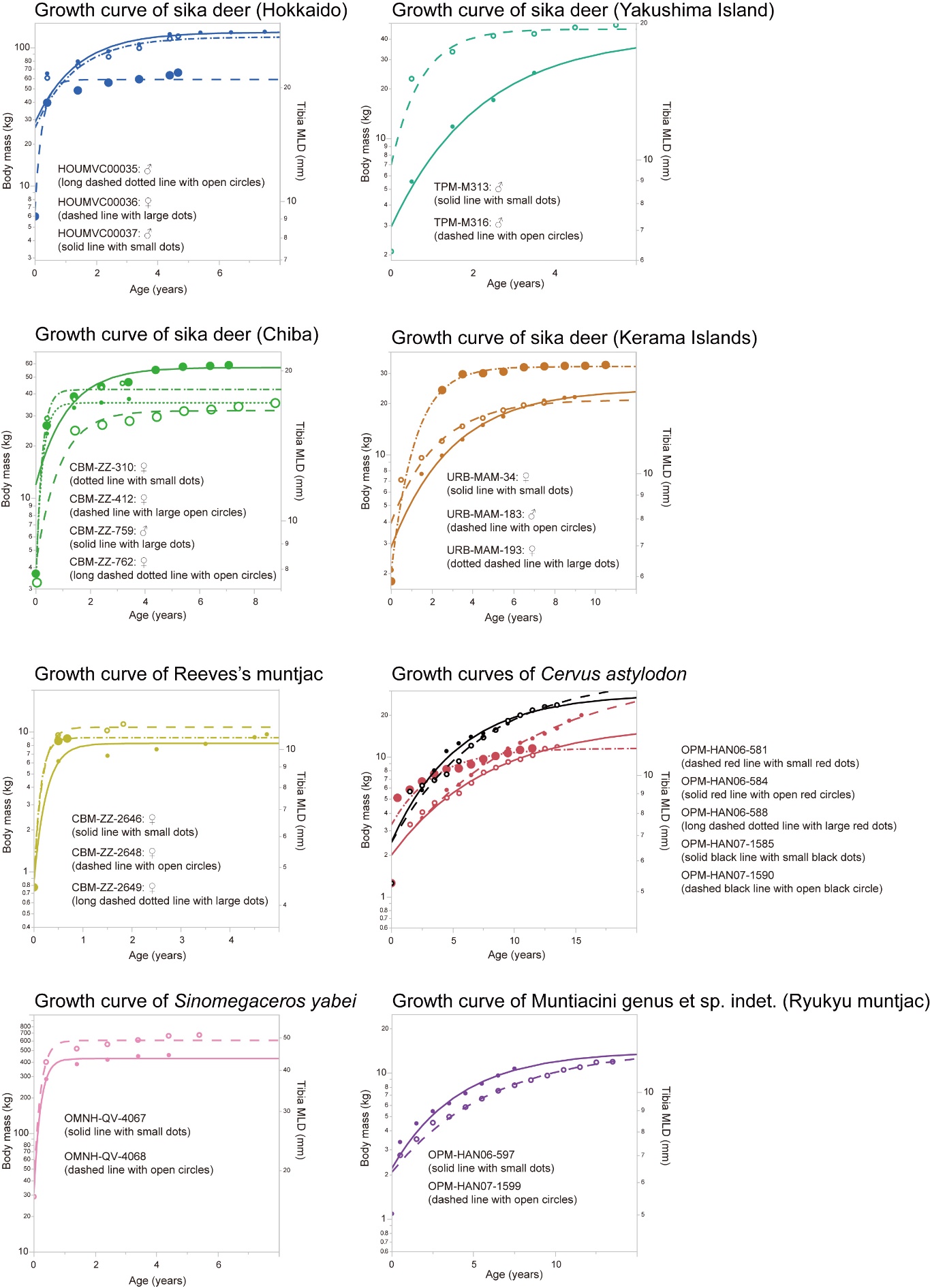

**Figure S9. Growth curves of body size estimates, i.e. the tibia mediolateral diameter (MLD) in the right axis and body mass estimated from the tibia MLD in left Y-axis, of eight cervid populations. The Gompertz curves are fitted and the parameters for which (*a*: asymptote; *b*: growth rate; and *c*: inflection point) are shown in Table S8.**

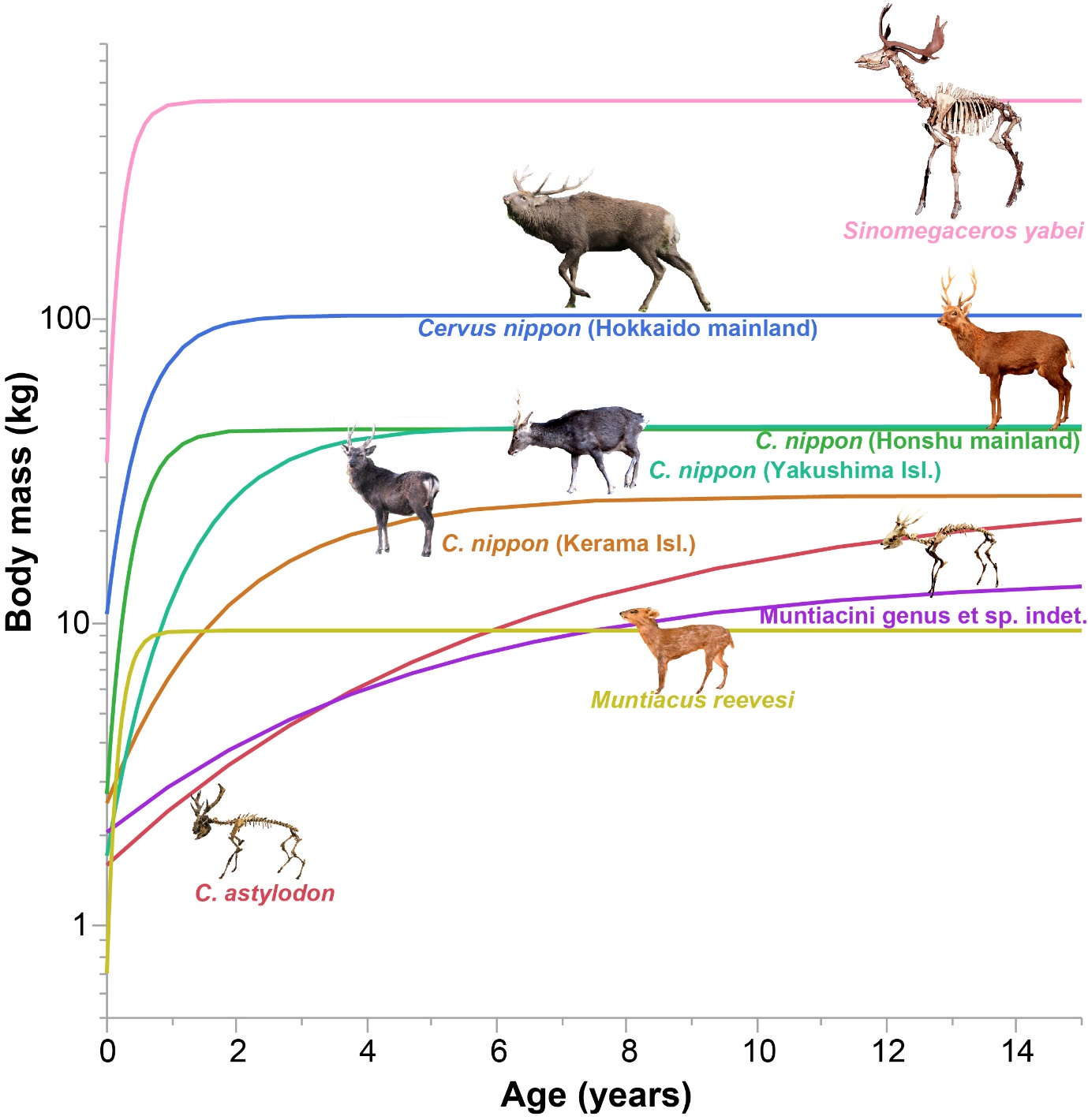

**Figure S10. Growth curves of insular cervids in Japan based on lines of arrested growth (LAGs) in the tibiae. Mainland cervids show rapid growth, attaining their maximum body size within 2 years, whereas two fossil Okinawa cervids exhibited an extremely slow growth rate, reaching their maximum body size at ca. 14 years for *Cervus astylodon* and 10 years for the Ryukyu muntjac (Muntiacini gen. et. sp. indet.). Sika deer (*Cervus nippon*) from Yakushima and Kerama Islands have an intermediate growth rate, attaining their maximum body size at around 5 years of age.**

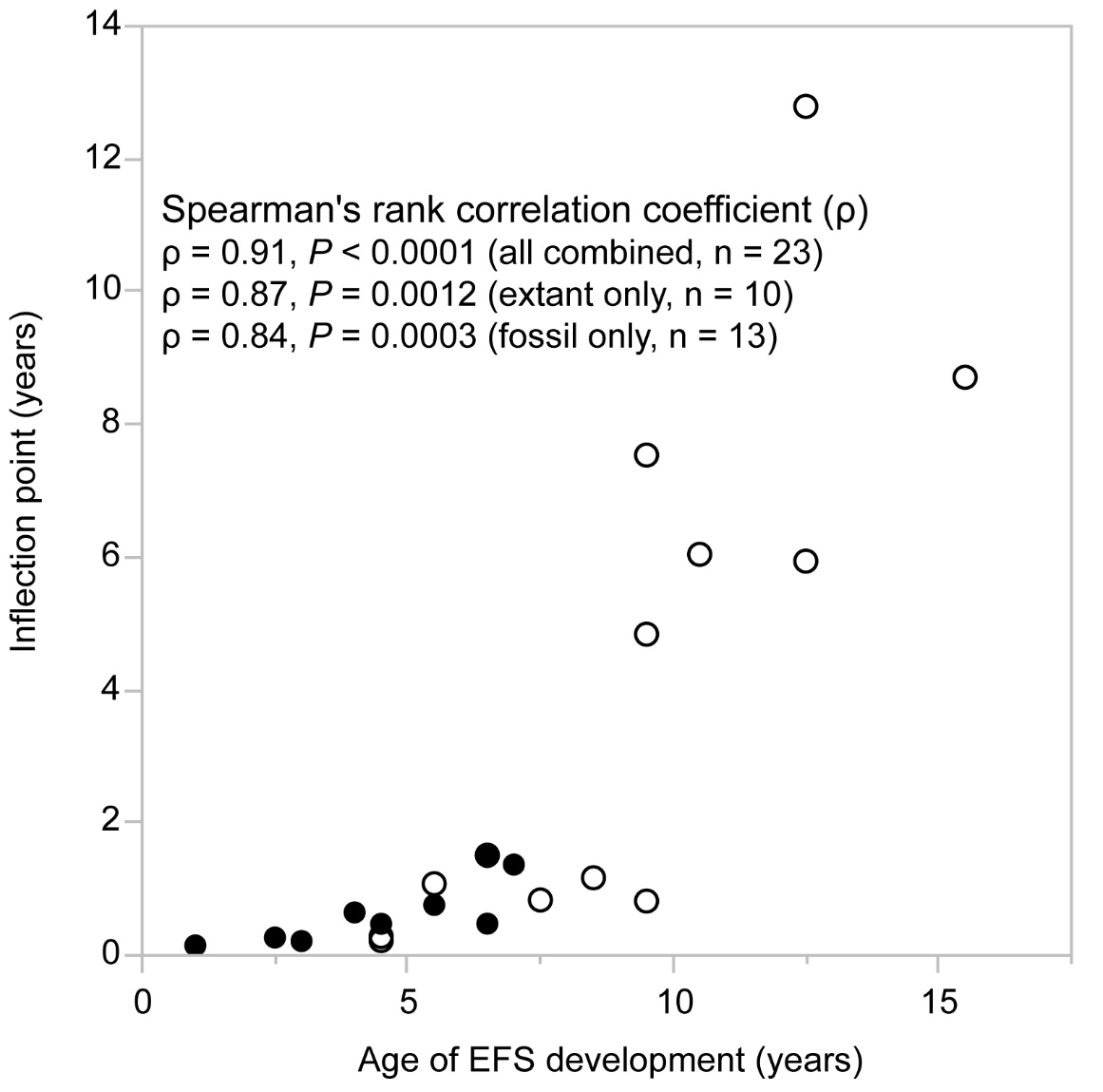

**Figure S11. Correlation between the inflection point, which was obtained from the fitted growth curve, and the age at which EFS development started. The values were obtained from Table S1 – S3 for the EFS developmental age and Table S8 for the inflection point. As for the extant specimens, the inflection point was averaged for femur and tibia growth models. The black dots represent extant samples, whereas the open circles being the fossil samples.**

**Supplementary Tables**

Table S1. Information about estimated body mass and age, epiphyseal closure and histological features in the femora of the fossil cervids.

| **Species** | **Collection no.** | **EBM**  **(kg) ^†^** | **GL**  **(mm)** | **Number of LAGs ^*^** | **Estimated age**  **(year)** | | **Epiphyseal fusion (p/d)** | | **Bone tissue type** | | **EFS ^**^** | **Degree of remodeling** |
| --- | --- | --- | --- | --- | --- | --- | --- | --- | --- | --- | --- | --- |
| ***Cervus astylodon*** | OPM-HAN07-1610 | 10.0 | 101.3 ^a^ | 6 | 6 | -/- | | PFB | | No | | Poor (only inner cortex) |
|  | OPM-HAN07-1612 | 11.2 | 104.3 ^a^ | 6 | 8 | -/Un | | PFB | | No | | Poor (only inner cortex) |
|  | OPM-HAN07-1604 | 11.9 | 112.1 ^a^ | 9 | 9 | Fg/- | | PFB | | No | | Poor (only inner cortex) |
|  | OPM-HAN06-151 | 17.9 | 124.1 ^a^ | 4 | 8 | -/Fd | | PFB | | No | | Extensive (PBTs are present only in the outer cortex) |
|  | OPM-HAN06-155 | 27.6 | 129.5 ^a^ | 11 (3) | 18 | -/- | | PFB | | Yes  (15 – 16 Y) | | Moderate (scattered SOs are present throughout the entire cortex) |
|  | OPM-HAN07-1603 | 20.4 | 130.4 ^a^ | 12 (3) | 15 | -/- | | PFB | | Yes  (12 – 13 Y) | | Moderate (scattered SOs are present throughout the entire cortex) |
| **Muntiacini gen. et sp. indet. (Ryukyu muntjac)** | OPM-HAN06-163 | 17.0 | 112.9 ^a^ | 11 (3) | 11 | Fd/- | | PFB | | Yes  (8 – 9 Y) | | Moderate (scattered SOs are present throughout the entire cortex) |
|  | OPM-HAN07-1618 | 15.1 | 121.4 ^a^ | 8 (2) | 8 | -/- | | PFB | | Yes  (6 – 7 Y) | | Moderate (scattered SOs are present throughout the entire cortex) |
|  | OPM-HAN07-1623 | 14.4 | 134.3 ^a^ | 9 (2) | 9 | -/- | | PFB | | Yes  (7 – 8 Y) | | Moderate (scattered SOs are present throughout the entire cortex) |
|  | OPM-HAN07-1616 | 22.1 | 144.4 ^a^ | 6 (1) | 6 | Fd/- | | PFB | | Yes  (5 – 6 Y) | | Moderate (scattered SOs are present throughout the entire cortex) |
| ***Sinomegaceros yabei*** | OMNH-QV-0362 | 397.6 | 391.7 ^a^ | 4 | 4 | Un/- | | FBL | | No | | Moderate (only posterior side of the cortex) |
|  | OMNH-QV-4062 | 959.2 | 465.0 | 7 (3) | 7 | Fd/- | | FBL | | Yes  (4 – 5 Y) | | Moderate (only posterior side of the cortex) |

Abbreviations: d, distal; EBM, estimated body mass; EFS, external fundamental system; FBL, fibro-lamellar bone tissue; Fd, fused; Fg, fusing; GL, greatest length of femur; LAG, line of arrested growth; p, proximal; PBT, primary bone tissue; PFB, parallel-fibered bone tissue; SO, secondary osteon; Un, unfused.

Collection acronyms: OPM, Okinawa Prefectural Museum and Art Museum, Okinawa, Japan; OMNH, Osaka Museum of Natural History, Osaka, Japan.

^†^ The estimated body mass (EBM) was obtained from the estimation equation based on femur mediolateral diameter. ^*^ The number of LAGs in EFS is shown in parenthesis. ^**^ The estimated age in years at which EFS started to develop is shown in parentheses. ^a^ The estimated greatest length of the bone based on measurements of the complete femora of other individuals. The number of LAGs in EFS is shown in parenthesis.

Table S2. Information about estimated body mass and age, epiphyseal closure and histological features in the tibiae of the fossil cervids.

| **Species** | **Collection no.** | **EBM**  **(kg) ^†^** | **GL**  **(mm)** | **Number of LAGs ^*^** | **Estimated age (year)** | **Epiphyseal fusion (p/d)** | **Bone tissue type** | **EFS ^**^** | **Degree of remodeling** |
| --- | --- | --- | --- | --- | --- | --- | --- | --- | --- |
| ***Cervus astylodon*** | OPM-HAN06-588 | 11.6 | 118.9 ^a^ | 11 (2) | 11 | -/Fd | PFB | Yes  (9 – 10 Y) | Moderate (scattered SOs are present throughout the entire cortex) |
|  | OPM-HAN06-584 | 11.9 | 122.5 ^a^ | 12 (3) | 13 | -/Fd | PFB | Yes  (10 – 11 Y) | Moderate (scattered SO s are present throughout the entire cortex) |
|  | OPM-HAN06-581 | 20.1 | 146.0 ^a^ | 13 (3) | 15 | -/Fd | FBL (from perimedullary region to 3^rd^ LAGs), PFB | Yes  (12 – 13 Y) | Moderate (scattered SOs are present throughout the entire cortex) |
|  | OPM-HAN07-1585 | 23.3 | 147.0 ^a^ | 10 (3) | 12 | -/- | PFB | Yes  (9 – 10 Y) | Moderate (scattered SOs are present throughout the entire cortex) |
|  | OPM-HAN07-1590 | 23.8 | 148.1 ^a^ | 12 (4) | 13 | -/- | PFB | Yes  (9 – 10 Y) | Moderate (scattered SOs are present throughout the entire cortex) |
| **Muntiacini gen. et sp. indet. (Ryukyu muntjac)** | OPM-HAN06-597 | 10.7 | 112.2 ^a^ | 7 (2) | 7 | -/Fd | PFB | Yes  (5 – 6 Y) | Moderate (scattered SOs are present throughout the entire cortex) |
|  | OPM-HAN07-1599 | 11.9 | 117.2 ^a^ | 13 (3) | 13 | -/- | PFB | Yes  (10 – 11 Y) | Moderate (scattered SOs are present throughout the entire cortex) |
| ***Sinomegaceros yabei*** | OMNH-QV-4067 | 458.6 | 406.7 ^a^ | 4 | 4 | -/- | FBL | No | Moderate (only anterior side of the cortex) |
|  | OMNH-QV-4068 | 678.6 | 444.9 ^a^ | 5 (1) | 5 | Fd/- | FBL | Yes  (4 – 5 Y) | Moderate (only anterior side of the cortex) |

Abbreviations and collection acronyms are shown in Table S1.

^†^ The estimated body mass (EBM) was obtained from the estimation equation based on tibia mediolateral diameter. ^*^ The number of LAGs in EFS is shown in parenthesis. ^**^ The estimated age in years at which EFS started to develop is shown in parentheses. ^a^ The estimated greatest length of the bone based on measurements of the complete femora of other individuals.

Table S3. Information about estimated body mass and age, epiphyseal closure and presence of EFS in the femora and tibiae of extant cervids.

|  |  |  |  | **GL (mm)** | |  | **Number of LAGs^*^** | | **Epiphyseal**  **fusion** | | **Bone tissue type** | **EFS** | **Degree of remodeling** | |
| --- | --- | --- | --- | --- | --- | --- | --- | --- | --- | --- | --- | --- | --- | --- |
| **Population** | **Collection no.** | **Sex** | **EBM**  **(kg) ^†^** | **Femur** | **Tibia** | **Age^‡^** | **Femur** | **Tibia** | **Femur**  **(p/d)** | **Tibia (p/d)** | **Femur**  **& tibia** | **Femur & tibia^**^** | **Femur** | **Tibia** |
| **Sika deer (*Cervus nippon*) (Hokkaido, mainland)** | HOUMVC  00039 | ♂ | 56.2 | 235 | 282 | 8M | 1 | 1 | Un/Un | Un/Un | FBL | No | Poor (only posterior side of the cortex) | Poor (only innermost cortex) |
|  | HOUMVC  00038 | ♀ | 38.0 | 218 | 270 | 8M | 1 | 1 | Un/Un | Un/Un | FBL | No | Poor (only posterior side of the cortex) | None |
|  | HOUMVC  00033 | ♂ | 94.5 | 284 | 337 | 1Y8M | 2 | 2 | Un/Un | Un/Fd | FBL | No | Poor (only posterior side of the cortex) | Extensive (only anterior side of the cortex) |
|  | HOUMVC  00034 | ♀ | 96.7 | 271 | 319 | 1Y9M | 2 | 2 | Un/Un | Un/Un | FBL | No | Moderate (only posterior side of the cortex) | None |
|  | HOUMVC  00035 | ♂ | 127.2 | 299 | 356 | 4Y8M | 5 | 5 | Fd/Un | Un/Fd | FBL | No | Extensive (only posterior side of the cortex) | Extensive (only anterior side of the cortex) |
|  | HOUMVC  00036 | ♀ | 85.8 | 243 | 295 | 4Y8M | 5 (2) | 5 (2) | Fd/Fd | Fd/Fd | FBL | Yes  (2 – 3Y) | Extensive (only posterior side of the cortex) | Poor (only innermost cortex) |
|  | HOUMVC  00037 | ♂ | 175.5^a^ | 300 | 346 | 7Y6M | 7 (3) | 7 (3) | Fd/Fd | Fd/Fd | FBL | Yes  (4 – 5Y) | Extensive (only posterior side of the cortex) | Poor (only innermost cortex) |
| **Sika deer (Honshu, mainland)** | CBM-ZZ-757 | NA | 3.5 | 104.3 | 128.1 | 0 D | 0 | 0 | Un/Un | Un/Un | WB | No | None | None |
|  | CBM-ZZ-671 | ♀ | 22.2 | 170.5^a^ | 202 | 6M | 1 | 1 | Un/- | Un/Un | FBL | No | Poor (only a few erosion cavities) | None |
| **Sika deer (Honshu, mainland)** | CBM-ZZ-218 | ♀ | 28.2 | 191.4 | 212.6 | 1Y2M | 1 | 1 | Un/Un | Un/Un | FBL | No | Poor (only a few erosion cavities) | Moderate (only anterior side of the cortex) |
|  | CBM-ZZ-761 | ♂ | 35.6 | 203.4 | 239.4 | 1Y2M | 1 | 1 | Un/Un | Un/Un | FBL | No | Poor (only a few erosion cavities) | Poor (only a few erosion cavities) |
|  | CBM-ZZ-310 | ♀ | 47.0 | 217.4 | 253 | 3Y5M | 2 (0) | 2 (0) | Fd/Fd | Fd/Fd | FBL | Yes  (3Y) | Extensive (only posterior side of the cortex) | Extensive (only anterior side of the cortex) |
|  | CBM-ZZ-762 | ♂ | 51.0 | 252 | 292.4 | 3Y2M | 3 | 3 | Fg/Un | Fg/Fd | FBL | No | Extensive (only posterior side of the cortex) | Moderate (scattered SOs are present throughout the entire cortex) |
|  | CBM-ZZ-759 | ♂ | 62.8 | 237.6 | 277.6 | 7Y1M | 7 (1) | 7 (1) | Fd/Fd | Fd/Fd | FBL | Yes  (6 – 7Y) | Extensive (only posterior side of the cortex) | Extensive (only anterior side of the cortex) |
|  | CBM-ZZ-412 | ♀ | 47.8 | 218.6 | 258.8 | 8Y9M | 7 (3) | 7 (3) | Fd/Fd | Fd/Fd | FBL | Yes  (5 – 6Y) | Extensive (only posterior side of the cortex) | Extensive (only anterior side of the cortex) |
| **Sika deer (Yakushima Island)** | TPM-M-312 | NA | 13.5 | 142^a^ | 160.8^a^ | 0Y | 0 | 0 | Un/Un | Un/Un | FBL | No | None | None |
| **Sika deer (Yakushima Island)** | TPM-M-314 | ♂ | 30.8 | 174.9^a^ | 213.2^a^ | 1Y6M | 1 | 1 | Un/Un | Un/Fd | FBL | No | Poor (only a few erosion cavities) | Poor (only a few erosion cavities) |
|  | TPM-M-313 | ♂ | 24.4 | 179 | 207.4 | 3Y6M | 3 | 3 | Un/Un | Un/Fd | FBL | No | Extensive (only posterior side of the cortex) | Extensive (only anterior side of the cortex) |
|  | TPM-M-316 | ♂ | 35.8 | 216.6 | 249.6 | 5Y6M | 5 (2) | 5 (1) | Fd/Fd | Fd/Fd | FBL | Yes  (3 – 5Y) | Extensive (only posterior side of the cortex) | Extensive (only anterior side of the cortex) |
| **Sika deer (Kerama Islands)** | URB-MAM-  80 | NA | 7.2 | 120.1 | 147.7 | 4M | 0 | 0 | Un/Un | Un/Un | FBL | No | None | None |
|  | URB-MAM-  211 | ♀ | 13.8 | 167.8 | 195 | 1Y | 1 | 1 | Un/Un | Un/Un | FBL | No | None | None |
|  | URB-MAM-  194 | ♂ | 18.4 | 168.3 | 198 | 1Y | 1 | 1 | Un/Un | Un/Un | FBL | No | None | None |
|  | URB-MAM-  34 | ♀ | 26.6 | 182.7 | 218 | NA | 8 (3) | 8 (3) | Fd/Fd | Fd/Fd | FBL, PFB | Yes | Extensive (only inner cortex) | Moderate (only inner cortex) |
|  | URB-MAM-  183 | ♂ | 20.8 | 184.3 | 215 | 7Y6M | 7 (0) | 7 (0) | Fd/Fd | Fd/Fd | FBL, PFB | Yes  (7Y) | Moderate (only inner cortex) | Extensive (only anterior side of the cortex) |
|  | URB-MAM-  55 | ♂ | 21.6 | 185.5 | 229 | 4Y6M | 7 | 7 | Un/Un | Un/Un | FBL, PFB | No | Extensive (only posterior side of the cortex) | Extensive (only anterior side of the cortex) |
| **Sika deer (Kerama Islands)** | URB-MAM-  193 | ♀ | 35.8 | 198.4 | 224 | 10Y6M | 8 (3) | 8 (5) | Fd/Fd | Fd/Fd | FBL, PFB | Yes  (5 – 8Y) | Extensive (only posterior side of the cortex) | Extensive (only anterior side of the cortex) |
| **Reeves’s muntjac (*Muntiacus reevesi*)** | CBM-ZZ-  4974 | NA | 0.8 | 34.3 | 40.6 | 0D | 0 | 0 | Un/Un | Un/Un | WB | No | None | None |
|  | CBM-ZZ-  2687 | ♂ | 4.9 ^b^ | 110.7 | 116.9 | 5M | 0 | 0 | Un/Un | Un/Un | FBL | No | Poor (scattered erosion cavities on posterior side of the cortex) | Poor (scattered erosion cavities on posterior side of the cortex) |
|  | CBM-ZZ-  2649 | ♀ | 6.8 ^b^ | 124.6 | 126.4 | 10M | 1 | 1 | Un/Un | Un/Fd | FBL | No | Poor (scattered erosion cavities on posterior side of the cortex) | Poor (scattered erosion cavities on posterior side of the cortex) |
|  | CBM-ZZ-  2648 | ♀ | 8.5 ^b^ | 127.5 | 135 | 1Y10M | 2 (0) | 2 (0) | Fd/Fd | Fd/Fd | FBL | Yes  (1Y) | Extensive (only posterior side of the cortex) | Extensive (only posterior side of the cortex) |
|  | CBM-ZZ-  2646 | ♀ | 7.2 ^b^ | 126.4 | 135 | 4Y9M | 5 (1) | 5 (2) | Fd/Fd | Fd/Fd | FBL | Yes  (2 – 4Y) | Extensive (only posterior side of the cortex) | Extensive (only posterior side of the cortex) |

Note that the presence of EFS is correlated with epiphyseal fusion, suggesting the somatic maturity of the animals (see also [11]).

D, days old; M, months old; NA, unknown; WB, woven bone; Y, years old. See Table S1 for explanations of other abbreviations.

Collection acronyms: CBM-ZZ, Natural History Museum and Institute, Chiba, Japan; TPM, Tochigi Prefectural Museum, Tochigi, Japan; HOUMVC, the Hokkaido University Museum, Sapporo, Japan; URB, University of the Ryukyus, Okinawa, Japan.

^†^ The estimated body mass (EBM) was obtained from the estimation equation based on femur mediolateral diameter. ^‡^ Age at death was determined through analyses of tooth eruption and the cementum annuli for sika deer and tooth eruption and the molar wear score for Reeves’s muntjac. ^*^ The number of LAGs in EFS is shown in parentheses. ^**^ The estimated age in years at which EFS started to develop is shown in parentheses. ^a^ The estimated greatest length of the bone based on measurements of the complete femur/tibiae of other individuals. ^b^ The actual body mass measurement.

**Table S4. List of the extant sika deer (*Cervus nippon*) samples that were used to check the validity of using lines of arrested growth (LAGs) to estimate age.**

| **Population** | **Collection no.** | **Sex** | **Age assessment method** | **Age at death** | **Date or season**  **of death (year/mm/dd)** | **No. of winter seasons** | **No. of LAGs** |
| --- | --- | --- | --- | --- | --- | --- | --- |
| **Sika deer**  **(Hokkaido mainland)** | HOUMVC  00039 | ♂ | Tooth eruption | 8M | 1977/2/20 | 1 | 1 |
|  | HOUMVC  00038 | ♀ | Tooth eruption | 8M | 1977/2/23 | 1 | 1 |
|  | HOUMVC  00033 | ♂ | Tooth eruption | 1Y8M | 1976/2/29 | 2 | 2 |
|  | HOUMVC  00034 | ♀ | Tooth eruption | 1Y9M | 1976/2/29 | 2 | 2 |
|  | HOUMVC  00035 | ♂ | Cementum annuli | 4Y8M | 1976/2/26 | 5 | 5 |
|  | HOUMVC  00036 | ♀ | Cementum annuli | 4Y8M | 1977/2/20 | 5 | 5 |
|  | HOUMVC  00037 | ♂ | Cementum annuli | 7Y6M | 1955/12/19 | 8 | 7 |
| **Sika deer**  **(Honshu mainland)** | CMB-ZZ-671 | ♀ | Tooth eruption | 6M | 1987/12/19 | 1 | 1 |
|  | CBM-ZZ-218 | ♀ | Tooth eruption | 1Y2M | 1986/8/9 | 1 | 1 |
|  | CBM-ZZ-761 | ♂ | Tooth eruption | 1Y2M | 1988/8/8 | 1 | 1 |
|  | CBM-ZZ-310 | ♀ | Cementum annuli | 3Y5M | 1986/11/19 | 3 | 3 |
|  | CBM-ZZ-762 | ♂ | Cementum annuli | 3Y2M | 1988/8/9 | 3 | 3 |
|  | CBM-ZZ-759 | ♂ | Cementum annuli | 7Y1M | 1988/7/4 | 7 | 7 |
|  | CBM-ZZ-412 | ♀ | Cementum annuli | 8Y9M | 1987/3/12 | 8 | 7 |
| **Sika deer**  **(Yakushima Island)** | TPM-M314 | ♂ | Tooth eruption | 1Y6M | Autumn–winter | 1 | 1 |
|  | TPM-M313 | ♂ | Cementum annuli | 3Y6M | Autumn–winter | 3 | 3 |
|  | TPM-M316 | ♂ | Cementum annuli | 5Y6M | Autumn–winter | 5 | 5 |
| **Sika deer**  **(Kerama Islands)** | URB-MAM-55 | ♂ | Cementum annuli | 4Y6M | Autumn–winter? | 4 | 7 |
|  | URB-MAM-183 | ♂ | Cementum annuli | 7Y6M | Autumn–winter? | 7 | 7 |
|  | URB-MAM-193 | ♀ | Cementum annuli | 10Y6M | Autumn–winter? | 11 | 8 |

Since LAGs form during winter, we compared the number of LAGs with the number of winter seasons a deer had experienced during its lifetime (See also Figure S5).

Abbreviations: M, months; Y, years.

**Table S5. Estimation of the number of lines of arrested growth (LAGs) that were lost through bone remodeling in fossil specimens.**

| **Species** | **Bone** | **Collection no.** | **No. observed**  **LAGs** | **Reference**  **specimen** | **Expansion of**  **medullary cavity** | **Estimated**  **LAGs lost** | **Estimated age at death (years)** |
| --- | --- | --- | --- | --- | --- | --- | --- |
| ***Cervus astylodon*** | Femur | OPM-HAN07-1610 | 6 | NA | NA | NA | 6 |
|  |  | OPM-HAN07-1612 | 6 | OPM-HAN07-1610 | Yes (2 LAGs) | 2 | 8 |
|  |  | OPM-HAN07-1604 | 9 | OPM-HAN07-1610 | No | 0 | 9 |
|  |  | OPM-HAN07-1603 | 12 | OPM-HAN07-1610 | Yes (3 LAGs) | 3 | 15 |
|  |  | OPM-HAN06-155 | 11 | OPM-HAN07-1610 | Yes (7 LAGs) | 7 | 18 |
|  |  | OPM-HAN06-151 | 4 | OPM-HAN07-1610 | Yes (4 LAGs) | 4 | 8 |
|  | Tibia | OPM-HAN06-588 | 11 | NA | NA | 0 | 11 |
|  |  | OPM-HAN06-584 | 12 | OPM-HAN06-588 | Yes (1 LAG) | 1 | 13 |
|  |  | OPM-HAN07-1585 | 10 | OPM-HAN06-588 | Yes (2 LAGs) | 2 | 12 |
|  |  | OPM-HAN06-581 | 13 | OPM-HAN06-588 | Yes (2 LAGs) | 2 | 15 |
|  |  | OPM-HAN07-1590 | 12 | OPM-HAN06-588 | Yes (1 LAG) | 1 | 13 |
| **Muntiacini gen. et. sp. indet.**  **(Ryukyu muntjac)** | Femur | OPM-HAN07-1618 | 8 | NA | NA | 0 | 8 |
|  |  | OPM-HAN07-1623 | 9 | OPM-HAN07-1618 | No | 0 | 9 |
|  |  | OPM-HAN07-1616 | 6 | OPM-HAN07-1618 | No | 0 | 6 |
|  |  | OPM-HAN06-163 | 11 | OPM-HAN07-1618 | No | 0 | 11 |
|  | Tibia | OPM-HAN06-597 | 7 | NA | NA | 0 | 7 |
|  |  | OPM-HAN07-1599 | 13 | OPM-HAN06-597 | No | 0 | 13 |
| ***Sinomegaceros yabei*** | Femur | OMNH-QV-0362 | 4 | NA | NA | 0 | 4 |
|  |  | OMNH-QV-4062 | 7 | OMNH-QV-0362 | No | 0 | 7 |
|  | Tibia | OMNH-QV-4067 | 4 | NA | NA | 0 | 4 |
|  |  | OMNH-QV-4068 | 5 | OMNH-QV-4067 | No | 0 | 5 |

**Table S6. Estimation of the neonatal body mass of the fossil cervids. Estimates were made using the regression equation of neonatal body mass against adult body mass for 121 extant artiodactyls (see also Figure S6).**

|  | **Estimated adult body mass (kg)** | | | **Estimated neonatal**  **body mass (kg)** |
| --- | --- | --- | --- | --- |
| **Species** | **Femur MLD** | **Tibia MLD** | **Average** |  |
| ***Cervus astylodon*** | 15.531 | 17.424 | 16.477 | 1.2588 |
| **Muntiacini gen. et. sp. indet.**  **(Ryukyu muntjac)** | 16.273 | 11.307 | 13.790 | 1.0905 |
| ***Sinomegaceros yabei*** | 959.175 | 678.566 | 818.870 | 29.378 |

MLD, mediolateral diameter.

**Table S7. Results of model selection for ontogenetic body mass change.**

| **Fitted model** | **Femur-based body mass data** | | | | |  | **Tibia-based body mass data** | | | | |
| --- | --- | --- | --- | --- | --- | --- | --- | --- | --- | --- | --- |
|  | **AICc** | **SSE** | **MSE** | **RMSE** | **R^2^** |  | **AICc** | **SSE** | **MSE** | **RMSE** | **R^2^** |
| **Gompertz (3 parameters)** | 2013.14 | 39254.37 | 276.44 | 16.63 | 0.9927 |  | 1787.19 | 23624.09 | 169.96 | 13.04 | 0.9910 |
| **Logistic (3 parameters)** | 2030.46 | 42470.00 | 299.08 | 17.29 | 0.9921 |  | 1795.43 | 24579.21 | 176.83 | 13.30 | 0.9906 |
| **Gompertz (4 parameters)** | 2135.49 | 33536.81 | 289.11 | 17.00 | 0.9938 |  | 2085.50 | 53881.62 | 464.50 | 21.55 | 0.9794 |
| **Logistic (4 parameters)** | 2338.73 | 84475.41 | 728.24 | 26.99 | 0.9844 |  | 2821.84 | 1857251.40 | 16010.79 | 126.53 | 0.2907 |

**Table S8. The parameter values of the fitted Gomperz curves, together with standard errors (S.E.), 95% confidence limits (C.L.), test statistics and P-values. Coefficients of determinations (R^2^) of the fitted Gompertz models are also provided.**

| **Population/Species** | **Collection no.** | **Bone** | **Parameter** | **Estimate** | **95%C.L.** | | **S.E.** | **Wald statistic** | **P-value**  **(χ^2^ test)** | **R^2^** |
| --- | --- | --- | --- | --- | --- | --- | --- | --- | --- | --- |
|  |  |  |  |  | **Lower** | **Upper** |  |  |  |  |
| **Sika deer**  **(Hokkaido, mainland)** | HOUMVC 00035 | F | a | 111.72 | 99.32 | 124.12 | 6.33 | 311.95 | <0.001 | 0.93 |
|  |  |  | b | 4.43 | 0.38 | 8.49 | 2.07 | 4.59 | 0.032 |  |
|  |  |  | c | 0.23 | 0.05 | 0.40 | 0.09 | 6.33 | 0.012 |  |
|  |  | T | a | 119.45 | 82.93 | 155.96 | 18.63 | 41.10 | <0.001 | 0.89 |
|  |  |  | b | 0.76 | -0.06 | 1.58 | 0.42 | 3.33 | 0.068 |  |
|  |  |  | c | 0.54 | -0.20 | 1.28 | 0.38 | 2.07 | 0.150 |  |
|  | HOUMVC 00036 | F | a | 80.27 | 68.40 | 92.13 | 6.05 | 175.77 | <0.001 | 0.92 |
|  |  |  | b | 1.28 | 0.25 | 2.31 | 0.53 | 5.94 | 0.015 |  |
|  |  |  | c | 0.35 | -0.02 | 0.72 | 0.19 | 3.43 | 0.064 |  |
|  |  | T | a | 58.68 | 52.94 | 64.42 | 2.93 | 401.47 | <0.001 | 0.93 |
|  |  |  | b | 4.23 | 1.20 | 7.26 | 1.55 | 7.47 | 0.006 |  |
|  |  |  | c | 0.19 | 0.04 | 0.33 | 0.07 | 6.19 | 0.013 |  |
|  | HOUMVC 00037 | F | a | 132.49 | 123.29 | 141.69 | 4.69 | 797.14 | <0.001 | 0.96 |
|  |  |  | b | 1.52 | 0.78 | 2.26 | 0.38 | 16.14 | <0.001 |  |
|  |  |  | c | 0.39 | 0.18 | 0.61 | 0.11 | 13.25 | <0.001 |  |
|  |  | T | a | 128.77 | 110.73 | 146.81 | 9.21 | 195.67 | <0.001 | 0.91 |
|  |  |  | b | 0.74 | 0.26 | 1.22 | 0.25 | 8.98 | 0.003 |  |
|  |  |  | c | 0.55 | 0.03 | 1.08 | 0.27 | 4.35 | 0.037 |  |
| **Sika deer**  **(Honshu, mainland)** | CBM-ZZ-310 | F | a | 44.68 | 41.99 | 47.38 | 1.38 | 1054.99 | <0.001 | 0.99 |
|  |  |  | b | 3.99 | 2.45 | 5.53 | 0.78 | 25.82 | <0.001 |  |
|  |  |  | c | 0.23 | 0.16 | 0.31 | 0.04 | 37.83 | <0.001 |  |
|  |  | T | a | 35.57 | 33.31 | 37.84 | 1.16 | 948.17 | <0.001 | 0.99 |
|  |  |  | b | 4.33 | 2.73 | 5.94 | 0.82 | 28.03 | <0.001 |  |
|  |  |  | c | 0.20 | 0.12 | 0.27 | 0.04 | 26.37 | <0.001 |  |
| **Sika deer**  **(Honshu, mainland)** | CBM-ZZ-412 | F | a | 45.32 | 39.96 | 50.69 | 2.74 | 274.21 | <0.001 | 0.94 |
|  |  |  | b | 0.59 | 0.29 | 0.89 | 0.15 | 14.61 | <0.001 |  |
|  |  |  | c | 0.94 | 0.37 | 1.51 | 0.29 | 10.57 | 0.001 |  |
|  |  | T | a | 32.34 | 30.04 | 34.65 | 1.18 | 757.09 | <0.001 | 0.94 |
|  |  |  | b | 1.16 | 0.63 | 1.69 | 0.27 | 18.40 | <0.001 |  |
|  |  |  | c | 0.58 | 0.20 | 0.95 | 0.19 | 9.24 | 0.002 |  |
|  | CBM-ZZ-759 | F | a | 55.63 | 50.69 | 60.58 | 2.52 | 486.74 | <0.001 | 0.94 |
|  |  |  | b | 1.20 | 0.54 | 1.85 | 0.33 | 12.85 | <0.001 |  |
|  |  |  | c | 0.44 | 0.14 | 0.74 | 0.15 | 8.38 | 0.004 |  |
|  |  | T | a | 57.06 | 50.82 | 63.29 | 3.18 | 321.72 | <0.001 | 0.93 |
|  |  |  | b | 0.87 | 0.38 | 1.36 | 0.25 | 12.03 | <0.001 |  |
|  |  |  | c | 0.51 | 0.11 | 0.91 | 0.20 | 6.22 | 0.013 |  |
|  | CBM-ZZ-762 | F | a | 45.74 | 39.22 | 52.26 | 3.33 | 188.89 | <0.001 | 0.96 |
|  |  |  | b | 3.24 | 0.27 | 6.21 | 1.52 | 4.56 | 0.033 |  |
|  |  |  | c | 0.25 | 0.06 | 0.43 | 0.09 | 6.93 | 0.008 |  |
|  |  | T | a | 47.85 | 42.10 | 53.60 | 2.93 | 266.18 | <0.001 | 0.97 |
|  |  |  | b | 4.62 | 1.38 | 7.86 | 1.65 | 7.79 | 0.005 |  |
|  |  |  | c | 0.20 | 0.06 | 0.34 | 0.07 | 7.44 | 0.006 |  |
| **Sika deer**  **(Yakushima Island)** | TMP-M-313 | F | a | 24.37 | 16.87 | 31.87 | 3.83 | 40.54 | <0.001 | 0.95 |
|  |  |  | b | 1.06 | 0.04 | 2.07 | 0.52 | 4.19 | 0.041 |  |
|  |  |  | c | 0.53 | 0.00 | 1.06 | 0.27 | 3.78 | 0.052 |  |
|  |  | T | a | 42.08 | 14.12 | 70.05 | 14.27 | 8.70 | 0.003 | 0.99 |
|  |  |  | b | 0.46 | 0.16 | 0.76 | 0.15 | 8.81 | 0.003 |  |
|  |  |  | c | 2.12 | 0.49 | 3.76 | 0.83 | 6.48 | 0.011 |  |
|  | TPM-M-316 | F | a | 36.23 | 31.60 | 40.86 | 2.36 | 235.16 | <0.001 | 0.98 |
|  |  |  | b | 0.80 | 0.45 | 1.16 | 0.18 | 19.96 | <0.001 |  |
|  |  |  | c | 0.83 | 0.48 | 1.17 | 0.18 | 22.21 | <0.001 |  |
| **Sika deer**  **(Yakushima Island)** | TPM-M-316 | T | a | 46.20 | 41.05 | 51.35 | 2.63 | 308.80 | <0.001 | 0.96 |
|  |  |  | b | 1.36 | 0.54 | 2.18 | 0.42 | 10.61 | 0.001 |  |
|  |  |  | c | 0.47 | 0.17 | 0.76 | 0.15 | 9.83 | 0.002 |  |
| **Sika deer**  **(Kerama Islands)** | URB-MAM-34 | F | a | 25.90 | 23.29 | 28.51 | 1.33 | 378.49 | <0.001 | 0.95 |
|  |  |  | b | 0.53 | 0.30 | 0.77 | 0.12 | 19.46 | <0.001 |  |
|  |  |  | c | 1.07 | 0.53 | 1.60 | 0.27 | 15.12 | <0.001 |  |
|  |  | T | a | 24.29 | 22.19 | 26.39 | 1.07 | 511.83 | <0.001 | 0.99 |
|  |  |  | b | 0.34 | 0.27 | 0.41 | 0.04 | 87.13 | <0.001 |  |
|  |  |  | c | 2.27 | 1.92 | 2.63 | 0.18 | 157.91 | <0.001 |  |
|  | URB-MAM-183 | F | a | 23.58 | 14.58 | 32.58 | 4.59 | 26.36 | <0.001 | 0.95 |
|  |  |  | b | 0.30 | 0.08 | 0.52 | 0.11 | 7.07 | 0.008 |  |
|  |  |  | c | 1.59 | 0.07 | 3.10 | 0.77 | 4.19 | 0.041 |  |
|  |  | T | a | 21.10 | 18.36 | 23.83 | 1.39 | 228.88 | <0.001 | 0.98 |
|  |  |  | b | 0.45 | 0.28 | 0.62 | 0.08 | 28.30 | <0.001 |  |
|  |  |  | c | 1.15 | 0.70 | 1.60 | 0.23 | 25.12 | <0.001 |  |
|  | URB-MAM-193 | F | a | 34.97 | 32.86 | 37.08 | 1.08 | 1053.64 | <0.001 | 0.98 |
|  |  |  | b | 0.46 | 0.35 | 0.58 | 0.06 | 60.93 | <0.001 |  |
|  |  |  | c | 1.77 | 1.39 | 2.16 | 0.20 | 82.43 | <0.001 |  |
|  |  | T | a | 32.95 | 32.24 | 33.65 | 0.36 | 8349.34 | <0.001 | 0.99 |
|  |  |  | b | 0.86 | 0.74 | 0.99 | 0.06 | 176.93 | <0.001 |  |
|  |  |  | c | 1.19 | 0.98 | 1.40 | 0.11 | 121.19 | <0.001 |  |
| **Reeves’s muntjac (*Muntiacus reevesi*)** | CBM-ZZ-264 | F | a | 8.38 | 7.45 | 9.32 | 0.48 | 309.26 | <0.001 | 0.92 |
|  |  |  | b | 3.67 | 0.77 | 6.57 | 1.48 | 6.13 | 0.013 |  |
|  |  |  | c | 0.22 | 0.02 | 0.42 | 0.10 | 4.66 | 0.031 |  |
|  |  | T | a | 8.31 | 7.28 | 9.34 | 0.53 | 249.35 | <0.001 | 0.90 |
|  |  |  | b | 3.98 | 0.53 | 7.43 | 1.76 | 5.12 | 0.024 |  |
|  |  |  | c | 0.21 | -0.01 | 0.43 | 0.11 | 3.44 | 0.064 |  |
| **Reeves’s muntjac (*Muntiacus reevesi*)** | CBM-ZZ-2648 | F | a | 7.47 | 6.90 | 8.05 | 0.29 | 643.00 | <0.001 | 0.99 |
|  |  |  | b | 6.06 | 3.30 | 8.82 | 1.41 | 18.53 | <0.001 |  |
|  |  |  | c | 0.14 | 0.05 | 0.22 | 0.04 | 10.27 | 0.001 |  |
|  |  | T | a | 10.82 | 9.69 | 11.95 | 0.57 | 354.55 | <0.001 | 0.99 |
|  |  |  | b | 6.02 | 2.65 | 9.40 | 1.72 | 12.22 | <0.001 |  |
|  |  |  | c | 0.16 | 0.04 | 0.28 | 0.06 | 6.82 | 0.009 |  |
|  | CBM-ZZ-2649 | F | a | 11.14 | – | – | – | – | – | 1.00 |
|  |  |  | b | 8.09 | – | – | – | – | – |  |
|  |  |  | c | 0.12 | – | – | – | – | – |  |
|  |  | T | a | 9.09 | – | – | – | – | – | 1.00 |
|  |  |  | b | 7.58 | – | – | – | – | – |  |
|  |  |  | c | 0.12 | – | – | – | – | – |  |
| ***Sinomegaceros yabei*** | OMNH-QV-0362 | F | a | 367.04 | 339.58 | 394.51 | 14.01 | 686.09 | <0.001 | 0.97 |
|  |  |  | b | 6.62 | 3.36 | 9.89 | 1.66 | 15.83 | <0.001 |  |
|  |  |  | c | 0.14 | 0.05 | 0.23 | 0.05 | 8.63 | 0.003 |  |
|  | OMNH-QV-4062 | F | a | 851.33 | 793.65 | 909.01 | 29.43 | 836.96 | <0.001 | 0.95 |
|  |  |  | b | 3.40 | 0.88 | 5.92 | 1.28 | 7.01 | 0.008 |  |
|  |  |  | c | 0.28 | 0.15 | 0.42 | 0.07 | 17.13 | <0.001 |  |
|  | OMNH-QV-4067 | T | a | 428.86 | 396.62 | 461.10 | 16.45 | 679.72 | <0.001 | 0.98 |
|  |  |  | b | 4.65 | 2.23 | 7.07 | 1.23 | 14.17 | <0.001 |  |
|  |  |  | c | 0.21 | 0.10 | 0.31 | 0.05 | 15.16 | <0.001 |  |
|  | OMNH-QV-4068 | T | a | 610.58 | 551.97 | 669.19 | 29.90 | 416.91 | <0.001 | 0.94 |
|  |  |  | b | 4.75 | 0.91 | 8.58 | 1.96 | 5.89 | 0.015 |  |
|  |  |  | c | 0.22 | 0.07 | 0.38 | 0.08 | 7.97 | 0.005 |  |
| ***Cervus astylodon*** | OPM-HAN06-151 | F | a | 29.54 | 21.16 | 37.92 | 4.28 | 47.71 | <0.001 | 1.00 |
|  |  |  | b | 0.22 | 0.15 | 0.29 | 0.04 | 37.71 | <0.001 |  |
|  |  |  | c | 5.38 | 3.96 | 6.79 | 0.72 | 55.43 | <0.001 |  |
| ***Cervus astylodon*** | OPM-HAN06-155 | F | a | 36.11 | 32.98 | 39.25 | 1.60 | 508.66 | <0.001 | 1.00 |
|  |  |  | b | 0.13 | 0.11 | 0.15 | 0.01 | 188.28 | <0.001 |  |
|  |  |  | c | 8.71 | 8.00 | 9.42 | 0.36 | 581.87 | <0.001 |  |
|  | OPM-HAN07-1603 | F | a | 25.11 | 23.66 | 26.56 | 0.74 | 1156.50 | <0.001 | 1.00 |
|  |  |  | b | 0.17 | 0.15 | 0.19 | 0.01 | 364.79 | <0.001 |  |
|  |  |  | c | 5.94 | 5.54 | 6.34 | 0.20 | 860.80 | <0.001 |  |
|  | OPM-HAN07-1604 | F | a | 16.26 | 9.51 | 23.02 | 3.45 | 22.24 | <0.001 | 0.97 |
|  |  |  | b | 0.19 | 0.08 | 0.30 | 0.06 | 10.98 | <0.001 |  |
|  |  |  | c | 3.22 | 0.72 | 5.72 | 1.28 | 6.36 | 0.012 |  |
|  | OPM-HAN07-1610 | F | a | 16.13 | 7.06 | 25.21 | 4.63 | 12.13 | <0.001 | 0.98 |
|  |  |  | b | 0.26 | 0.11 | 0.40 | 0.08 | 11.45 | <0.001 |  |
|  |  |  | c | 3.69 | 1.22 | 6.17 | 1.26 | 8.56 | 0.003 |  |
|  | OPM-HAN07-1612 | F | a | 17.16 | 12.44 | 21.87 | 2.41 | 50.87 | <0.001 | 0.99 |
|  |  |  | b | 0.21 | 0.14 | 0.27 | 0.03 | 37.12 | <0.001 |  |
|  |  |  | c | 4.52 | 3.04 | 6.00 | 0.75 | 35.77 | <0.001 |  |
|  | OPM-HAN06-581 | T | a | 42.94 | 27.18 | 58.70 | 8.04 | 28.51 | <0.001 | 0.99 |
|  |  |  | b | 0.09 | 0.06 | 0.11 | 0.01 | 48.13 | <0.001 |  |
|  |  |  | c | 12.79 | 8.55 | 17.03 | 2.16 | 34.93 | <0.001 |  |
|  | OPM-HAN06-584 | T | a | 17.42 | 12.97 | 21.87 | 2.27 | 58.89 | <0.001 | 0.99 |
|  |  |  | b | 0.13 | 0.09 | 0.17 | 0.02 | 38.46 | <0.001 |  |
|  |  |  | c | 6.04 | 3.80 | 8.28 | 1.14 | 27.98 | <0.001 |  |
|  | OPM-HAN06-588 | T | a | 11.67 | 9.77 | 13.56 | 0.97 | 145.63 | <0.001 | 0.92 |
|  |  |  | b | 0.29 | 0.13 | 0.44 | 0.08 | 12.80 | <0.001 |  |
|  |  |  | c | 0.82 | -0.03 | 1.66 | 0.43 | 3.61 | 0.057 |  |
|  | OPM-HAN07-1585 | T | a | 28.55 | 21.70 | 35.40 | 3.50 | 66.73 | <0.001 | 0.98 |
|  |  |  | b | 0.18 | 0.11 | 0.26 | 0.04 | 26.07 | <0.001 |  |
|  |  |  | c | 4.84 | 3.33 | 6.34 | 0.77 | 39.62 | <0.001 |  |
| ***Cervus astylodon*** | OPM-HAN07-1590 | T | a | 38.47 | 26.58 | 50.37 | 6.07 | 40.18 | <0.001 | 0.99 |
|  |  |  | b | 0.13 | 0.09 | 0.18 | 0.02 | 34.31 | <0.001 |  |
|  |  |  | c | 7.54 | 4.95 | 10.12 | 1.32 | 32.72 | <0.001 |  |
| **Muntiacini gen. et sp. indet. (Ryukyu muntjac)** | OPM-HAN06-163 | F | a | 16.66 | 13.95 | 19.37 | 1.38 | 145.07 | <0.001 | 0.93 |
|  |  |  | b | 0.29 | 0.14 | 0.45 | 0.08 | 14.20 | <0.001 |  |
|  |  |  | c | 1.17 | 0.33 | 2.01 | 0.43 | 7.46 | 0.006 |  |
|  | OPM-HAN07-1616 | F | a | 22.78 | 16.11 | 29.44 | 3.40 | 44.80 | <0.001 | 0.93 |
|  |  |  | b | 0.47 | 0.11 | 0.83 | 0.18 | 6.67 | 0.010 |  |
|  |  |  | c | 1.08 | 0.18 | 1.97 | 0.46 | 5.57 | 0.018 |  |
|  | OPM-HAN07-1618 | F | a | 16.62 | 11.90 | 21.33 | 2.40 | 47.77 | <0.001 | 0.94 |
|  |  |  | b | 0.31 | 0.11 | 0.51 | 0.10 | 9.27 | 0.002 |  |
|  |  |  | c | 1.51 | 0.36 | 2.66 | 0.59 | 6.58 | 0.010 |  |
|  | OPM-HAN07-1623 | F | a | 14.00 | 11.67 | 16.34 | 1.19 | 138.38 | <0.001 | 0.92 |
|  |  |  | b | 0.39 | 0.16 | 0.62 | 0.12 | 11.05 | <0.001 |  |
|  |  |  | c | 0.83 | 0.10 | 1.57 | 0.38 | 4.90 | 0.027 |  |
|  | OPM-HAN06-597 | T | a | 14.03 | 7.49 | 20.58 | 3.34 | 17.65 | <0.001 | 0.97 |
|  |  |  | b | 0.24 | 0.08 | 0.41 | 0.08 | 8.49 | 0.004 |  |
|  |  |  | c | 2.50 | 0.29 | 4.72 | 1.13 | 4.90 | 0.027 |  |
|  | OPM-HAN07-1599 | T | a | 14.75 | 12.66 | 16.83 | 1.06 | 191.79 | <0.001 | 0.99 |
|  |  |  | b | 0.16 | 0.13 | 0.20 | 0.02 | 69.63 | <0.001 |  |
|  |  |  | c | 4.06 | 3.03 | 5.08 | 0.52 | 59.93 | <0.001 |  |

**Table S9. Statistical comparisons of the growth curve parameters (growth rate and inflection point) among eight extant and fossil cervids using one-way ANOVA. Since bone type (femur or tibia) did not have a statistically significant effect on the growth curve parameters (see SI Appendix text), the femur and tibia data were combined.**

|  |  | **Growth rate** | | | | | | | |  | **Inflection point** | | | | | | | |
| --- | --- | --- | --- | --- | --- | --- | --- | --- | --- | --- | --- | --- | --- | --- | --- | --- | --- | --- |
|  |  |  |  |  | **95% CI** | | **Statistical**  **tests** | | |  |  |  |  | **95% CI** | | | **Statistical**  **tests** | |
| **Population/species** |  | **n** | **Mean** | **S.E.** | **Lower** | **Upper** |  |  |  |  | **n** | **Mean** | **S.E.** | **Lower** | | **Upper** |  |  |
| **Reeves’s muntjac (*Muntiacus reevesi*)** |  | 3 | 5.901 | 0.543 | 4.788 | 7.013 | A |  |  |  | 3 | 0.161 | 1.129 | −2.152 | 2.474 | | A |  |
| ***Sinomegaceros yabei*** |  | 4 | 4.854 | 0.470 | 3.891 | 5.818 | A |  |  |  | 4 | 0.213 | 0.978 | −1.790 | 2.216 | | A |  |
| **Sika deer (*Cervus nippon*) (Honshu mainland)** |  | 4 | 2.499 | 0.470 | 1.536 | 3.462 |  | B |  |  | 4 | 0.418 | 0.978 | −1.585 | 2.422 | | A |  |
| **Sika deer (Hokkaido mainland)** |  | 3 | 2.161 | 0.543 | 1.049 | 3.273 |  | B | C |  | 3 | 0.375 | 1.129 | −1.938 | 2.688 | | A |  |
| **Sika deer (Yakushima Island)** |  | 2 | 0.921 | 0.665 | −0.441 | 2.283 |  | B | C |  | 2 | 0.986 | 1.383 | −1.847 | 3.819 | | A | B |
| **Sika deer (Kerama Islands)** |  | 3 | 0.490 | 0.543 | −0.622 | 1.603 |  | B | C |  | 3 | 1.506 | 1.129 | −0.807 | 3.819 | | A |  |
| **Ryukyu muntjac (Muntiacini gen. et. sp. indet.)** |  | 6 | 0.312 | 0.384 | −0.474 | 1.098 |  |  | C |  | 6 | 1.858 | 0.799 | 0.222 | 3.494 | | A |  |
| ***Cervus astylodon*** |  | 11 | 0.181 | 0.284 | −0.400 | 0.761 |  |  | C |  | 11 | 5.771 | 0.590 | 4.563 | 6.979 | |  | B |

CI, confidence interval; SE, standard error. Taxa with different letters in the “Statistical tests” columns are significantly different (Tukey–Kramer method, *P* < 0.05).

**Table S10. Statistical comparisons of the growth curve parameters (growth rate and inflection point) among eight extant and fossil cervids using two-way ANOVA with bone type and species/populations being the explanatory variables. Least square means were calculated, and the difference was tested by Tukey-Kramer method.**

|  |  | **Growth rate** | | | | | | | | |  | | **Inflection point** | | | | | | | | | | | | | | |
| --- | --- | --- | --- | --- | --- | --- | --- | --- | --- | --- | --- | --- | --- | --- | --- | --- | --- | --- | --- | --- | --- | --- | --- | --- | --- | --- | --- |
|  |  |  |  |  | **95% CI** | | **Statistical**  **tests** | | | |  | |  | | |  | | |  | | | **95% CI** | | | | **Statistical**  **tests** | |
| **Population/species** |  | **n** | **Mean** | **S.E.** | **Lower** | **Upper** |  |  |  |  |  | **n** | | | **Mean** | | | **S.E.** | | | **Lower** | | | **Upper** | |  |  |
| **Reeves’s muntjac (*Muntiacus reevesi*)** |  | 6 | 5.901 | 0.477 | 4.938 | 6.863 | A |  |  |  |  | | | 6 | | | 0.161 | | | 0.650 | | | −1.151 | | 1.473 | A |  |
| ***Sinomegaceros yabei*** |  | 4 | 4.854 | 0.584 | 3.675 | 6.034 | A |  |  |  |  | | | 4 | | | 0.213 | | | 0.796 | | | −1.394 | | 1.820 | A |  |
| **Sika deer (*Cervus nippon*) (Honshu mainland)** |  | 8 | 2.499 | 0.413 | 1.665 | 3.333 |  | B |  |  |  | | | 8 | | | 0.418 | | | 0.563 | | | −0.718 | | 1.555 | A |  |
| **Sika deer (Hokkaido mainland)** |  | 6 | 2.161 | 0.477 | 1.198 | 3.124 |  | B | C |  |  | | | 6 | | | 0.375 | | | 0.650 | | | −0.937 | | 1.687 | A |  |
| **Sika deer (Yakushima Island)** |  | 4 | 0.921 | 0.584 | −0.258 | 2.100 |  | B | C | D |  | | | 4 | | | 1.506 | | | 0.650 | | | 0.194 | | 2.818 | A |  |
| **Sika deer (Kerama Islands)** |  | 6 | 0.490 | 0.477 | −0.472 | 1.453 |  |  | C | D |  | | | 6 | | | 0.986 | | | 0.796 | | | −0.621 | | 2.593 | A |  |
| **Ryukyu muntjac (Muntiacini gen. et. sp. indet.)** |  | 6 | 0.307 | 0.480 | −0.662 | 1.276 |  |  | C | D |  | | | 6 | | | 1.947 | | | 0.654 | | | 0.626 | | 3.268 | A |  |
| ***Cervus astylodon*** |  | 11 | 0.179 | 0.353 | −0.532 | 0.891 |  |  |  | D |  | | | 11 | | | 5.795 | | | 0.481 | | | 4.825 | | 6.765 |  | B |

CI, confidence interval; SE, standard error. Taxa with different letters in the “Statistical tests” columns are significantly different (Tukey–Kramer method, *P* < 0.05).

**Table S11. Life tables for three extant sika deer (*Cervus nippon*) populations based on age data from randomly culled individuals.**

| **Population** | **Age**  **(years)** | **Age (% of maximum observed age)** | **Sampled fx** | **Fx (Probit smoothing)** | **lx** | **lx*1000** |
| --- | --- | --- | --- | --- | --- | --- |
| **Sika deer (Hokkaido mainland)** | 2 | 10 | 450 | 409 | 1.000 | **1000** |
|  | 3 | 15 | 242 | 236 | 0.577 | **577** |
|  | 4 | 20 | 113 | 141 | 0.345 | **345** |
|  | 5 | 25 | 63 | 88 | 0.214 | **214** |
|  | 6 | 30 | 46 | 56 | 0.138 | **138** |
|  | 7 | 35 | 34 | 37 | 0.091 | **91** |
|  | 8 | 40 | 37 | 25 | 0.062 | **62** |
|  | 9 | 45 | 24 | 18 | 0.043 | **43** |
|  | 10 | 50 | 14 | 13 | 0.031 | **31** |
|  | 11 | 55 | 15 | 9 | 0.022 | **22** |
|  | 12 | 60 | 6 | 7 | 0.016 | **16** |
|  | 13 | 65 | 6 | 5 | 0.012 | **12** |
|  | 14 | 70 | 3 | 4 | 0.009 | **9** |
|  | 15 | 75 | 2 | 3 | 0.007 | **7** |
|  | 16 | 80 | 2 | 2 | 0.005 | **5** |
|  | 17 | 85 | 2 | 2 | 0.004 | **4** |
|  | 18 | 90 | 0 | 1 | 0.003 | **3** |
|  | 19 | 95 | 1 | 1 | 0.003 | **3** |
|  |  |  | Total: 1060 |  |  |  |
| **Sika deer (Honshu mainland)** | 2 | 11 | 132 | 126 | 1.000 | **1000** |
|  | 3 | 17 | 108 | 108 | 0.853 | **853** |
|  | 4 | 22 | 78 | 89 | 0.708 | **708** |
|  | 5 | 28 | 70 | 72 | 0.571 | **571** |
|  | 6 | 33 | 51 | 56 | 0.448 | **448** |
|  | 7 | 39 | 42 | 43 | 0.342 | **342** |
|  | 8 | 44 | 34 | 32 | 0.254 | **254** |
|  | 9 | 50 | 24 | 23 | 0.184 | **184** |
|  | 10 | 56 | 25 | 16 | 0.130 | **130** |
|  | 11 | 61 | 14 | 11 | 0.089 | **89** |
|  | 12 | 67 | 7 | 8 | 0.060 | **60** |
|  | 13 | 72 | 7 | 5 | 0.040 | **40** |
|  | 14 | 78 | 1 | 3 | 0.026 | **26** |
|  | 15 | 83 | 0 | 2 | 0.016 | **16** |
|  | 16 | 89 | 0 | 1 | 0.010 | **10** |
|  | 17 | 94 | 1 | 1 | 0.006 | **6** |
|  |  |  | Total: 594 |  |  |  |
| **Sika deer (Yakushima Island)** | 2 | 11 | 18 | 18 | 1.000 | **1000** |
|  | 3 | 17 | 16 | 14 | 0.776 | **776** |
|  | 4 | 22 | 10 | 11 | 0.595 | **595** |
|  | 5 | 28 | 6 | 8 | 0.457 | **457** |
|  | 6 | 33 | 8 | 6 | 0.354 | **354** |
|  | 7 | 39 | 4 | 5 | 0.277 | **277** |
|  | 8 | 44 | 3 | 4 | 0.219 | **219** |
|  | 9 | 50 | 0 | 3 | 0.175 | **175** |
|  | 10 | 56 | 3 | 3 | 0.141 | **141** |
|  | 11 | 61 | 2 | 2 | 0.114 | **114** |
|  | 12 | 67 | 2 | 2 | 0.093 | **93** |
|  | 13 | 72 | 1 | 1 | 0.077 | **77** |
|  | 14 | 78 | 0 | 1 | 0.064 | **64** |
|  | 15 | 83 | 0 | 1 | 0.053 | **53** |
|  | 16 | 89 | 0 | 1 | 0.045 | **45** |
|  | 17 | 94 | 1 | 1 | 0.038 | **38** |
|  |  |  | Total: 74 |  |  |  |

**Table S12. Life tables for extant sika deer from Kinkazan Island, Reeves’s muntjac (*Muntiacus reevesi*) and two fossil insular deer based on age data from individuals that died naturally.**

| **Population** | **Age (years)** | **Age (% of maximum observed age)** | **Sampled fx** | **dx** | **lx** | **lx (Probit smoothing)** | **lx*1000** |
| --- | --- | --- | --- | --- | --- | --- | --- |
| ***Cervus nippon* (Kinkazan Isl.)** | 2 | 9 | 15 | 0.0568 | 1.000 | 0.975 | **1000** |
|  | 3 | 14 | 8 | 0.0303 | 0.943 | 0.955 | **980** |
|  | 4 | 18 | 12 | 0.0455 | 0.913 | 0.921 | **945** |
|  | 5 | 23 | 23 | 0.0871 | 0.867 | 0.865 | **888** |
|  | 6 | 27 | 29 | 0.1098 | 0.780 | 0.779 | **799** |
|  | 7 | 32 | 40 | 0.1515 | 0.670 | 0.660 | **677** |
|  | 8 | 36 | 43 | 0.1629 | 0.519 | 0.516 | **529** |
|  | 9 | 41 | 32 | 0.1212 | 0.356 | 0.369 | **379** |
|  | 10 | 45 | 23 | 0.0871 | 0.235 | 0.243 | **249** |
|  | 11 | 50 | 14 | 0.0530 | 0.148 | 0.150 | **154** |
|  | 12 | 55 | 8 | 0.0303 | 0.095 | 0.088 | **91** |
|  | 13 | 59 | 8 | 0.0303 | 0.064 | 0.051 | **52** |
|  | 14 | 64 | 2 | 0.0076 | 0.034 | 0.028 | **29** |
|  | 15 | 68 | 2 | 0.0076 | 0.027 | 0.016 | **16** |
|  | 16 | 73 | 1 | 0.0038 | 0.019 | 0.009 | **9** |
|  | 17 | 77 | 0 | 0.0000 | 0.015 | 0.005 | **5** |
|  | 18 | 82 | 2 | 0.0076 | 0.015 | 0.003 | **3** |
|  | 19 | 86 | 1 | 0.0038 | 0.008 | 0.001 | **1** |
|  | 20 | 91 | 0 | 0.0000 | 0.004 | 0.001 | **1** |
|  | 21 | 95 | 1 | 0.0038 | 0.004 | 0.000 | **0** |
|  |  |  | Total: 264 |  |  |  |  |
| ***Cervus astylodon*** | 2 | 8 | 3 | 0.067 | 1.000 | 0.952 | **1000** |
|  | 3 | 12 | 0 | 0.000 | 0.933 | 0.938 | **986** |
|  | 4 | 15 | 0 | 0.000 | 0.933 | 0.923 | **970** |
|  | 5 | 19 | 2 | 0.044 | 0.933 | 0.907 | **953** |
|  | 6 | 23 | 1 | 0.022 | 0.889 | 0.889 | **934** |
|  | 7 | 27 | 3 | 0.067 | 0.867 | 0.870 | **914** |
|  | 8 | 31 | 0 | 0.000 | 0.800 | 0.849 | **893** |
|  | 9 | 35 | 0 | 0.000 | 0.800 | 0.827 | **869** |
|  | 10 | 38 | 3 | 0.067 | 0.800 | 0.803 | **843** |
|  | 11 | 42 | 2 | 0.044 | 0.733 | 0.776 | **815** |
|  | 12 | 46 | 0 | 0.000 | 0.689 | 0.747 | **785** |
|  | 13 | 50 | 0 | 0.000 | 0.689 | 0.716 | **752** |
|  | 14 | 54 | 1 | 0.022 | 0.689 | 0.682 | **717** |
|  | 15 | 58 | 0 | 0.000 | 0.667 | 0.645 | **678** |
|  | 16 | 62 | 2 | 0.044 | 0.667 | 0.605 | **636** |
|  | 17 | 65 | 2 | 0.044 | 0.622 | 0.562 | **591** |
|  | 18 | 69 | 4 | 0.089 | 0.578 | 0.515 | **541** |
|  | 19 | 73 | 5 | 0.111 | 0.489 | 0.464 | **487** |
|  | 20 | 77 | 3 | 0.067 | 0.378 | 0.408 | **429** |
|  | 21 | 81 | 2 | 0.044 | 0.311 | 0.348 | **366** |
|  | 22 | 85 | 4 | 0.089 | 0.267 | 0.283 | **297** |
|  | 23 | 88 | 3 | 0.067 | 0.178 | 0.212 | **223** |
|  | 24 | 92 | 3 | 0.067 | 0.111 | 0.135 | **142** |
|  | 25 | 96 | 2 | 0.044 | 0.044 | 0.051 | **54** |
|  |  |  | Total: 45 |  |  |  |  |
| **Muntiacini gen. et sp. indet. (Ryukyu muntjac)** | 0 | 0 | 5 | 0.077 | 1.000 | 0.996 | **1000** |
|  | 1 | 9 | 2 | 0.031 | 0.923 | 0.958 | **961** |
|  | 2 | 18 | 6 | 0.092 | 0.892 | 0.912 | **915** |
|  | 3 | 27 | 1 | 0.015 | 0.800 | 0.858 | **861** |
|  | 4 | 36 | 0 | 0.000 | 0.785 | 0.794 | **797** |
|  | 5 | 45 | 2 | 0.031 | 0.785 | 0.719 | **722** |
|  | 6 | 55 | 3 | 0.046 | 0.754 | 0.630 | **632** |
|  | 7 | 64 | 27 | 0.415 | 0.708 | 0.525 | **527** |
|  | 8 | 73 | 16 | 0.246 | 0.292 | 0.400 | **402** |
|  | 9 | 82 | 2 | 0.031 | 0.046 | 0.254 | **254** |
|  | 10 | 91 | 1 | 0.015 | 0.015 | 0.080 | **80** |
|  |  |  | Total: 65 |  |  |  |  |
| **Extant Reeves’s muntjac** | 0 | 0 | 12 | 0.143 | 1.000 | 0.920 | **1000** |
|  | 1 | 6 | 21 | 0.250 | 0.857 | 0.798 | **868** |
|  | 2 | 13 | 9 | 0.107 | 0.607 | 0.684 | **744** |
|  | 3 | 19 | 7 | 0.083 | 0.500 | 0.578 | **629** |
|  | 4 | 25 | 3 | 0.036 | 0.417 | 0.482 | **524** |
|  | 5 | 31 | 8 | 0.095 | 0.381 | 0.395 | **429** |
|  | 6 | 38 | 2 | 0.024 | 0.286 | 0.318 | **346** |
|  | 7 | 44 | 3 | 0.036 | 0.262 | 0.251 | **273** |
|  | 8 | 50 | 4 | 0.048 | 0.226 | 0.194 | **211** |
|  | 9 | 56 | 5 | 0.060 | 0.179 | 0.147 | **160** |
|  | 10 | 63 | 1 | 0.012 | 0.119 | 0.109 | **118** |
|  | 11 | 69 | 0 | 0.000 | 0.107 | 0.078 | **85** |
|  | 12 | 75 | 1 | 0.012 | 0.107 | 0.054 | **59** |
|  | 13 | 81 | 2 | 0.024 | 0.095 | 0.037 | **40** |
|  | 14 | 88 | 2 | 0.024 | 0.071 | 0.024 | **26** |
|  | 15 | 94 | 4 | 0.048 | 0.048 | 0.015 | **16** |
|  |  |  | Total: 84 |  |  |  |  |
